# Affinity-enhanced peptides delivered by mRNA–LNPs inhibit influenza A virus replication by disrupting the PA-PB1 interaction

**DOI:** 10.64898/2026.07.29.740963

**Authors:** Alberto Florez Prada, Chloé Sturmach, Nabila Laroui, Nicolas Pautrieux, Camille Bourgneuf, Philippe Mas, Remy Lartia, Didier Boturyn, Wim P. Burmeister, Chantal Pichon, Nadia Naffakh, Catherine Isel, Darren J. Hart

## Abstract

Seasonal influenza causes up to 650,000 deaths annually and remains a persistent pandemic threat due to zoonotic strains crossing the species barrier. Current antivirals, which target neuraminidase or the viral polymerase, have limited efficacy and rapidly select for resistant variants, highlighting the urgent need for new therapeutic strategies. The heterotrimeric influenza polymerase (FluPol), comprising PA, PB1, and PB2 subunits, is essential for viral replication and harbors virus-specific protein-protein interfaces that are potential drug targets. We focused on the highly conserved PA-PB1 interface, where the N-terminal peptide of PB1 binds the PA C-terminal domain with high affinity. Disrupting this interaction abrogates polymerase function, halting viral replication. Using phage display, we identified PB1-derived peptides with enhanced affinity for PA and characterized their binding via biophysical methods and X-ray crystallography. Lead peptides efficiently disrupted the PB1-PA interaction and inhibited polymerase activity in cell-based assays. To address peptide delivery challenges, we expressed these inhibitors intracellularly from synthetic mRNA formulated in lipid nanoparticles, achieving robust inhibition of viral replication in cultured cells. This work establishes intracellularly expressed peptide inhibitors as a viable antiviral strategy and provides a generalizable framework for targeting essential protein-protein interactions of influenza and other RNA viruses.

## Introduction

Influenza is a leading cause of viral respiratory disease worldwide, infecting about 10% of adults and 30% of children annually. Seasonal influenza causes an estimated 290–650,000 deaths annually, with 70–85% fatalities occurring in older adults. Among children under 5 years of age, an estimated 28-111,000 deaths occur each year, with 99% of them in low- and middle-income countries (1). Vaccination is the primary preventive strategy, but uptake is limited, efficacy decreases in the elderly, and vaccines must be regularly updated to account for the antigenic drift. Mismatches with circulating strains, as seen in the 2025–2026 season, can significantly reduce vaccine effectiveness (2). In addition, animal influenza A viruses (IAVs) from novel antigenic subtypes can cross the species barrier, e.g., through genetic reassortment, become capable of sustained human-to-human transmission and cause pandemics (3). Effective anti-influenza drugs are needed to complement vaccination, for example, to treat high risk groups prophylactically, to treat critically ill patients, or to counter novel pandemic strains until a matched vaccine becomes available.

Current antiviral drugs target the sialidase activity of the surface glycoprotein neuraminidase (e.g. oseltamivir) and the endonuclease activity of the PA subunit of the viral RNA-dependent RNA polymerase (RdRp, FluPol, e.g. baloxavir) (4). Recent meta-analyses confirm that in hospitalized or high-risk patients, available antivirals offer reductions, although modest, in disease duration, with uncertain impacts on mortality (5). However, a strong limitation is that they must be administered shortly after the onset of symptoms to be effective, and the emergence of resistant variants is common. For example, 7% of baloxavir-treated out-patients in the 2024 Japanese influenza season exhibited the I38T/M resistance substitution (6), while oseltamivir-resistant H275Y variants are detected sporadically every winter (7), and circulated in a sustained manner during the 2007-2008 season (8). Thus, there is a critical need for new therapies with novel mechanisms of action to complement existing drugs (5) and may also open the way to combination therapies to limit resistance issues (9).

The heterotrimeric influenza polymerase, comprising the PA, PB1, and PB2 subunits, is essential for viral gene transcription and genome replication and has no homolog in host cells, making it an attractive antiviral target (10). A number of high-resolution structures of FluPol have been determined, providing key insights into its structure–function relationships (11). In its transcriptionally active state, FluPol adopts a characteristic U-shaped architecture, with two opposing protrusions corresponding to the PA N-terminal endonuclease domain (12, 13) and the PB2 cap-binding domain (14). The central body of the heterotrimer is formed by the catalytic core of PB1, which exhibits the canonical right-handed polymerase fold. The base of the U is constituted by the large C-terminal domain of PA (15, 16). Structural integrity of the complex is maintained through two major hydrophobic inter-subunit interfaces: one between the C-terminus of PB1 and the N-terminus of PB2 (17), and the other between the C-terminal domain of PA and the N-terminus of PB1 (15, 16), which together organize the subunits in a head-to-tail arrangement. Assembly of FluPol proceeds via PA–PB1 heterodimer formation in the cytoplasm, followed by nuclear import and association with PB2 in the nucleus to form the mature heterotrimer. The PA-PB1 heterodimer assembly proceeds via a conserved, highaffinity interaction between a short PB1 N-terminal 3_10_ helix corresponding to residues 1–15 (PB1_1–15_) and the PA C-terminal domain (PA_Cter_, residues 257 to 716), that has been structurally characterized at high resolution (Fig. 1) (15, 16, 18). The interaction of the PB1 peptide is mediated by the PA_Cter_ hydrophobic core, comprising an LLFL motif (PB1 residues 7–10), but stable complex formation also requires the terminal flanking residues. In particular, D2/V3/N4 contribute N-terminal contacts with PA, and residues around the C-terminal side of the motif, especially V12 (with additional contacts involving P13/A14), further stabilize the interface; accordingly, the PB1 core motif alone is not sufficient, and stable PA binding requires a peptide extending to PB1 residues 2–12.

**Figure 1.**
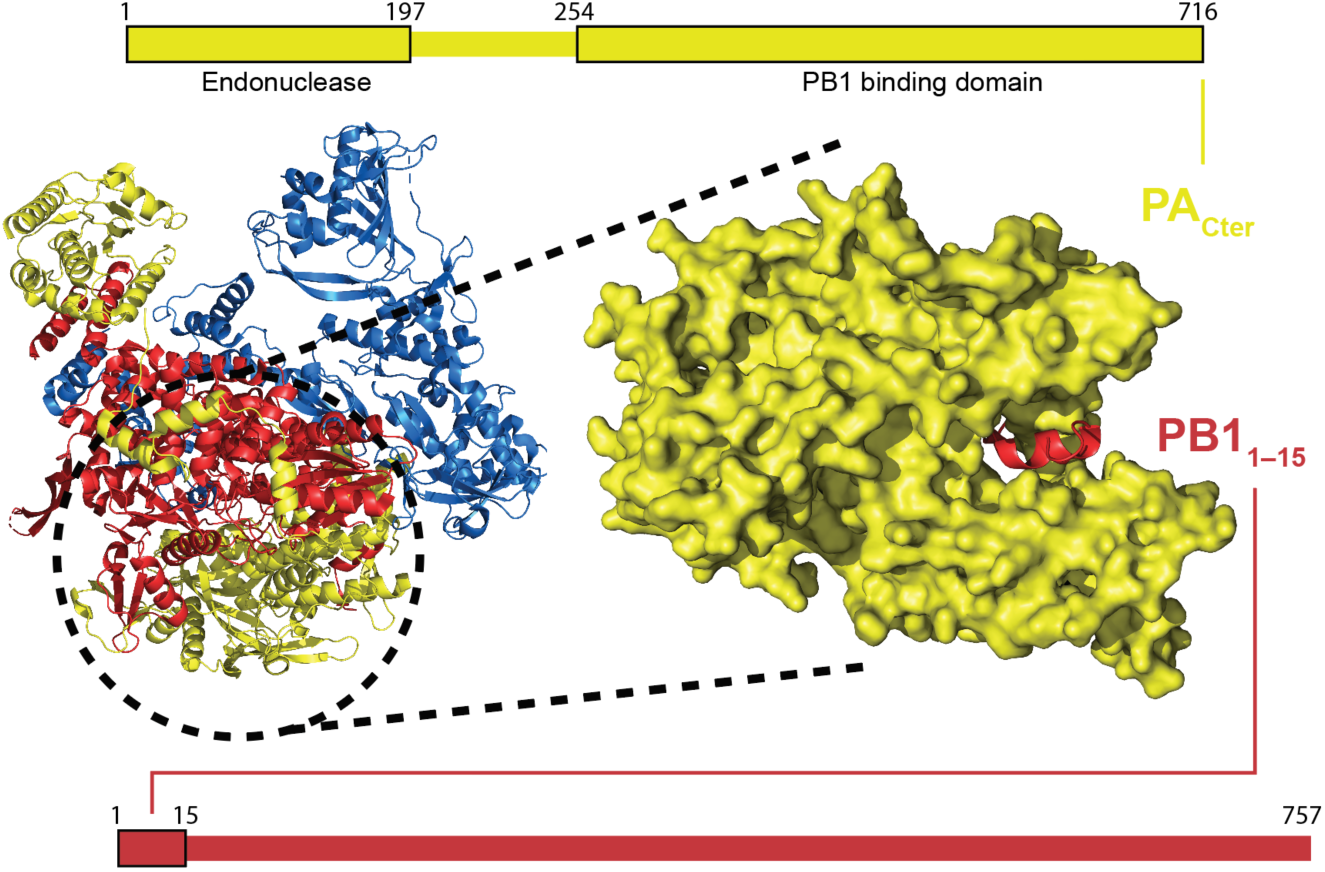
Domain architecture and structural basis of the PA–PB1 interaction in influenza polymerase. PA and PB1 domains are shown schematically (top and bottom), indicating the PA endonuclease domain (residues 1–197), the PA PB1-binding domain (residues 254–716), and the PB1 N-terminal region (residues 1-15) involved in PA binding. Left, ribbon representation of the influenza polymerase complex with PA colored yellow, PB1 red, and PB2 blue (PDB 4WSB); the circled region highlights the PA–PB1 interface. Right, zoomed surface view of the PA C-terminal domain bound to the PB1 N-terminal peptide (PDB 2ZNL), highlighting the interaction site.

Disrupting this interface abolishes FluPol activity, halting viral replication. Several studies have demonstrated that PB1_1–15_-derived peptides and small molecules can bind PA_Cter_ in vitro. Cell-based mini-replicon and infection assays confirm that these molecules successfully inhibit FluPol activity and reduce viral replication (19–21).

In general, protein-protein interactions (PPIs), often mediated by short linear motifs (SLiMs), are attractive drug targets because they rely on compact interaction surfaces, often defined by a few key affinity- and specificity-defining residues, yet these residues control essential functions (22, 23). PPI inhibition is often a domain of small molecules, which benefit from oral bioavailability and efficient intracellular delivery. However, PPIs can also be targeted by small peptides. Extracellular peptide antivirals (e.g., the HIV fusion inhibitor enfuvirtide) have reached the clinic but, in the context of respiratory viruses, such approaches increasingly compete with highly potent monoclonal antibodies, such as clesrovimab and nirsevimab, recently approved monoclonal antibodies against RSV (24, 25). Targeting intracellular PPIs with peptides remains challenging due to their short half-lives and limited membrane permeability (26, 27). Although chemical strategies such as macrocyclization and helix-stapling can improve stability and cellular uptake, robust intracellular peptide inhibitors remain uncommon (28, 29).

To bypass cell penetration challenges, we used an alternative approach: intracellular expression of therapeutic peptides from synthetic mRNA formulated as lipid nanoparticles (LNPs) (30, 31). These non-genome-integrating mRNA-LNP systems enable transient expression of polypeptides directly in the cytosol, following cellular uptake (31). Optimized LNP formulations for pulmonary or systemic delivery have achieved robust protein expression and favorable safety profiles in preclinical models (30, 31).

Here, we present a new strategy to inhibit IAV replication by targeting the PA-PB1 interface of FluPol. We first screened almost a billion random PB1_1–15_-derived peptides using phage display to maximize the affinity of this region for PA. We then characterized their binding modes using biophysical methods and X-ray crystallography and verified the ability of lead peptides to disrupt the PB1-PA dimer and inhibit FluPol activity in cell-based assays. Finally, we implemented a mRNA-LNP strategy for intracellular expression of peptide-carrier fusions. This approach addresses longstanding barriers in peptide vectorization and provides a practical framework for evaluating intracellular peptide drugs more generally.

## Results

### Phage display-guided optimization of PB1-derived peptides for high affinity PA binding

Guided by structural information on the IAV PA–PB1 interface (PDB 2ZNL (16)), we constructed two phage-display libraries focusing on the PA-contacting residues of the PB1_1–15_ N-terminal helix. In one M13 pIII-fused library, defined as Fully Randomized (FR), seven PA-contacting residues flanking the LLFL core (residues 1, 2, 4, 6, 11, 12 and 14) were fully randomized using NNS codons, where N = A/T/G/C and S = G/C (Fig. 1A). In a second library, defined as Rationally Randomized (RR), residue 6 was randomized with an NNS codon and seven hydrophobic positions (residues 1, 3, 7, 8, 9, 10, 12) were partially randomized using DKS codons, that are biased towards hydrophobic residues, where D = A/G/T, K = G/T and S = G/C, to permit controlled variation of the peptide core region, favoring hydrophobic identities and eliminating all stop codons (Fig. 1B). In an initial library selection experiment, both FR and RR libraries comprised about 2 × 10^5^ unique transformants. These were subjected to three rounds of panning against biotinylated PA C-terminal domain of influenza A strain WSN (A/WSN/1933 H1N1; WSN-PA_Cter_) using streptavidin-coated magnetic beads and short washes, which led to the identification of six peptide candidates after Sanger sequencing of ELISA-positive randomly selected colonies (32). We then optimized our workflow, producing substantially larger libraries of 7.7 × 10⁸ transformants for FR and 7.1 × 10⁸ for RR, and applying an increasingly stringent off-rate selection (33) over three rounds of panning. This was achieved by incubating phages with up to 5,000-fold excess of free WSN-PA_Cter_ in solution. We then used next-generation sequencing (NGS) to characterize selected binders. The total number of good quality translated sequence reads (encoding peptides) were18,091 for FR and 14,782 for RR. The diversity was greater in the RR library, with 4,408 unique sequences identified, compared with 1,886 in the FR library (Supp Table 1). Abundance distributions were more skewed in FR, where the top 18 sequences accounted for 54% of the reads (Figure 1A), than in RR, where the top 10 sequences accounted for 31% (Fig. 1B).

**Table 1.**
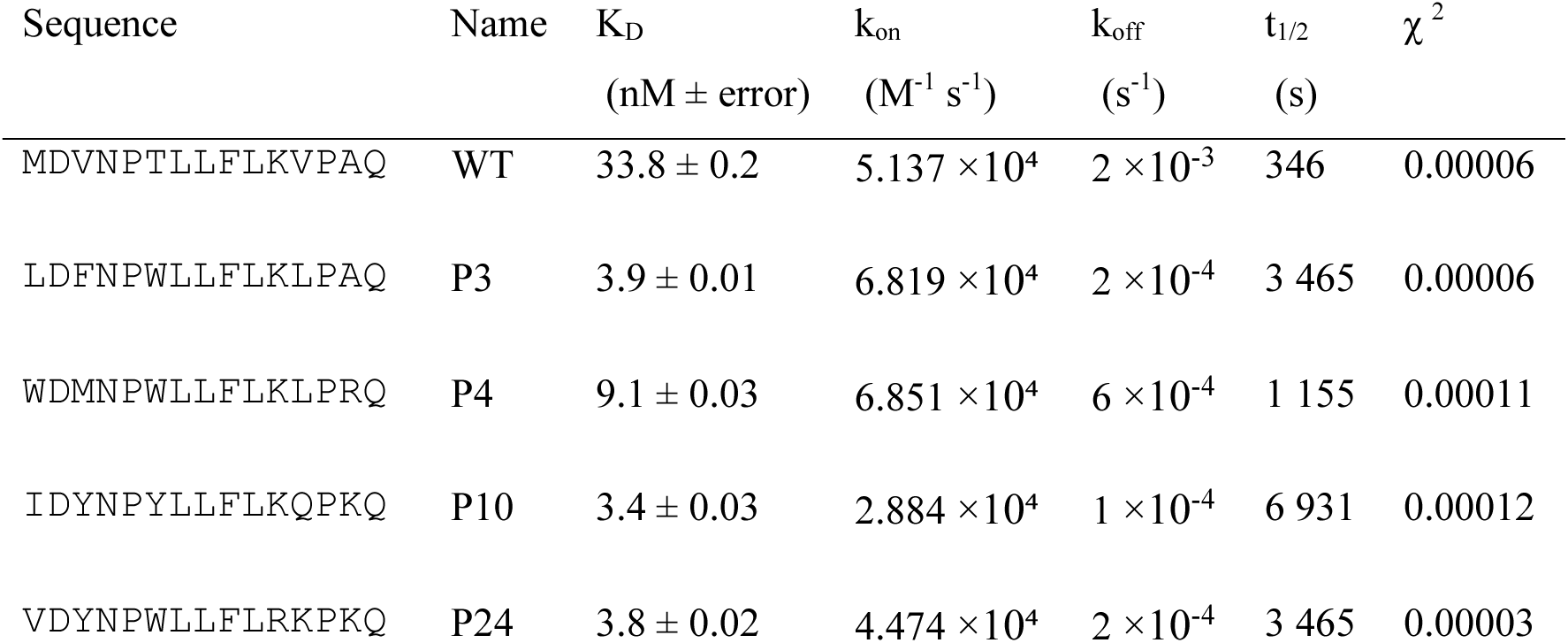
Kinetic parameters of WSN-PA_Cter_–peptide interactions. Dissociation constants (K_D_), association (k_on_) and dissociation (k_off_) rate constants, and t_1/2_ are reported with fit error estimates (χ^2^). The results were derived from a BLI run using 5 increasing WSN-PA_Cter_ concentrations.

Sequence logo representations of clones with >1% frequency revealed distinct trends: FR favored Tyr or Ile/Val at the N terminus and charged residues near the C terminus (Fig. 1C), whereas the RR library exhibited less positional bias, with hydrophobic residues such as Val, Leu, or Met enriched at the N terminus (Fig. 1D). Based on frequency and clustering by Hamming distance, 19 peptides (Supp. Table 2) were prioritized for further investigation, and a consensus peptide, P24, was designed by combining conserved N-ter, central and C-ter regions from the 19 selected peptides.

### Biophysical characterization of high-affinity peptides reveals enhanced binding and stability

Binding of the 19 selected peptides to PA_Cter_ was ranked by bio-layer interferometry (BLI). Raw BLI sensorgrams and global kinetic fits used to analyze the BLI data are shown in Supplementary Figs. 1 and 2, with processed sensorgrams and parameters shown in Fig. 3A and Table 1, respectively. The two lead highest affinity binders, hereby referred to as P3, P4, P10, and P24, were then re-evaluated against the wild-type PB1_1–15_ peptide (WT) and a scrambled control peptide (SCR; same amino acid composition as WT in random order) in a detailed BLI kinetic study.

**Figure 2.**
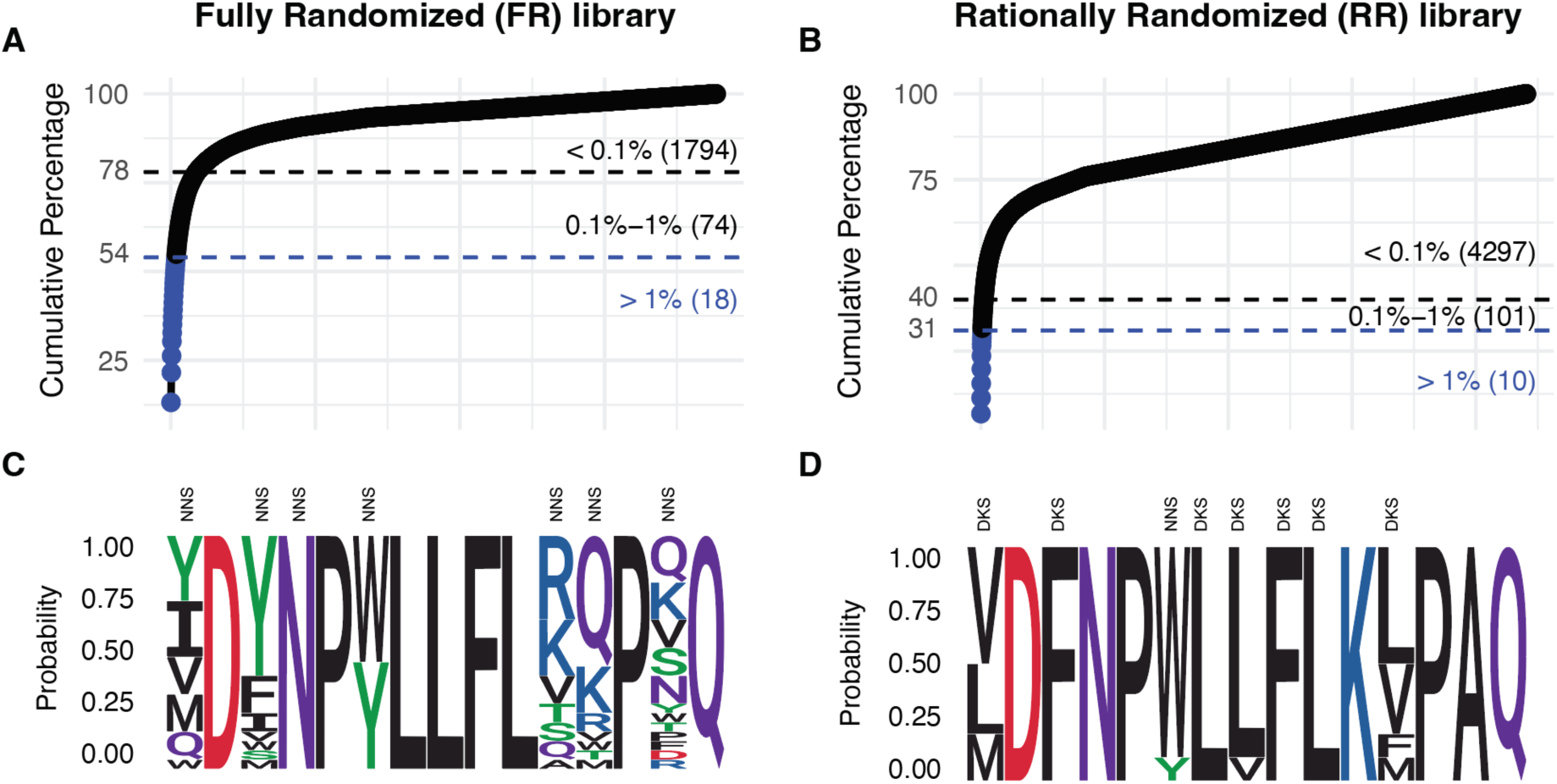
Sequence diversity of two phage display libraries targeting the PA–PB1 interaction of influenza A virus FluPol. **(A-B)**. Cumulative-percentage plots showing the frequency of unique peptide sequences in the fully randomized (FR) and rationally randomized (RR) libraries, with thresholds for >1%, 0.1–1%, and <0.1% abundance separated by black and blue lines; the FR library (A) contains 18 sequences at >1%, 74 at 0.1–1%, and 1,794 at <0.1%, the RR library (B) includes 10 sequences at >1%, 101 at 0.1–1%, and 4,297 at <0.1%. **(C-D)**. Amino acid distributions of unique sequences with >1% frequency (sequence logos), with degenerate codon types (NNS or DKS) indicated above randomized positions.

**Figure 3.**
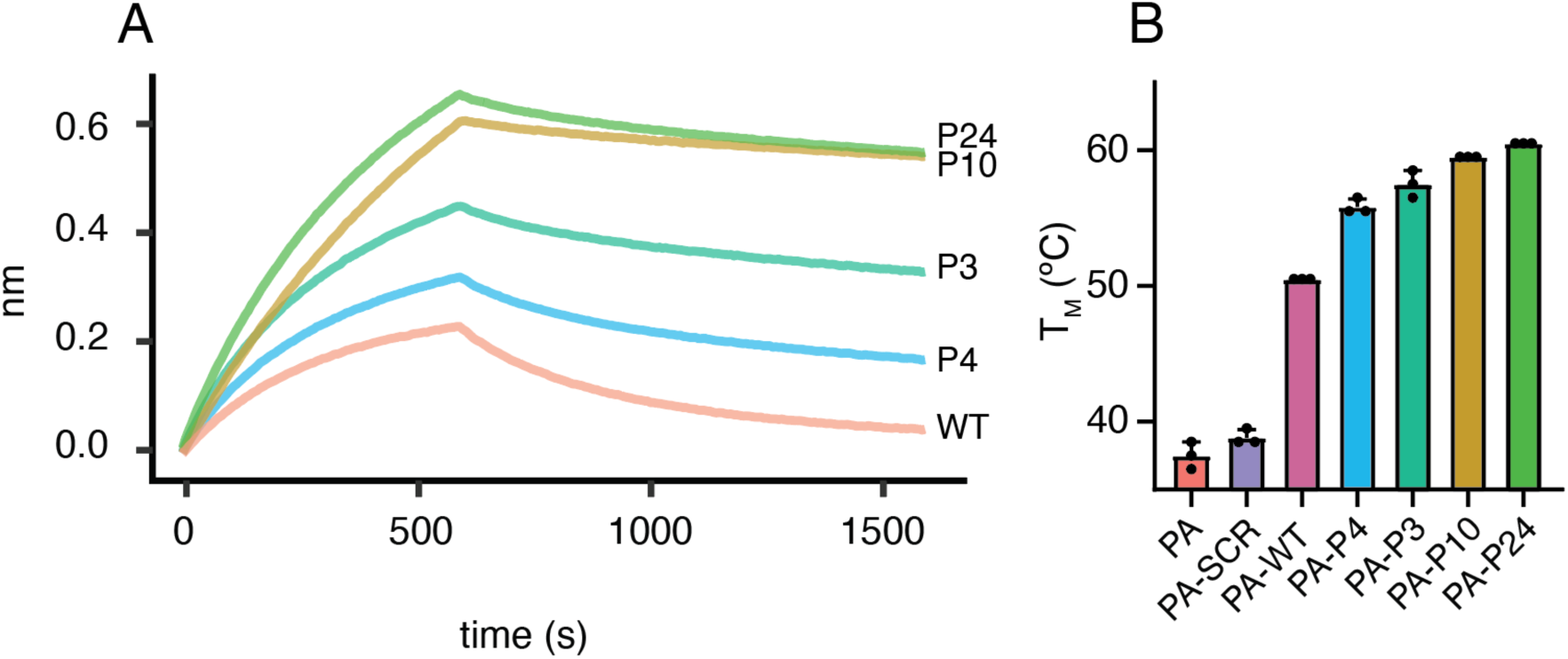
Biophysical characterization of selected peptides. **(A)** Representative **BLI** sensorgrams showing binding of WT and selected variants (P3, P4, P10 and P24) to WSN-PA_Cter_ at 60 nM. **(B)** Thermal shift assay showing the T_M_ of WSN-PA_Cter_ alone (PA), or in complex with scrambled control peptide (PA–SCR), wild-type PB1_1–15_ peptide (PA–WT), or selected variants (PA–P3, PA–P4, PA–P10, PA–P24). Each point represents a technical triplicate.

Peptide P4 bound with a K_D_ of 9.1 nM, representing a 3.7-fold improvement over WT at 33.8 nM, while P3, P10 and P24 bound more tightly, with K_D_ of 3.9, 3.4 and 3.8 nM respectively, corresponding to a ∼10-fold gain in affinity. Kinetic analyses revealed that the enhanced affinities were primarily due to slower dissociation rates. Whereas the WT exhibited a dissociation rate expressed as a half-life of 346 s, P3, P4, P10 and P24 displayed markedly longer half-lives (t_1/2_) of 3,465 s, 1,155 s, 6,931 s and 3,465 s, respectively. Association rates were similar or moderately higher than WT for P3 and P4 (k_on_ 5.1×10^4^ M^-1^s^-1^ vs ∼6.8×10^4^ M^-1^s^-1^) while P10 and P24 bound with slightly lower k_on_ values (2.9×10^4^ and 4.5×10^4^ M^-1^s^-1^, respectively).

Thermal shift assays (TSA) confirmed increased stability of the PA_Cter_–peptide complexes (Fig. 3B). PA_Cter_ alone or with the non-binding SCR control exhibited a melting temperature (T_M_) of ∼40 °C. Addition of WT raised the T_M_ by ∼10 °C, while addition of P3, P4, P10 or P24 increased the T_M_ by ∼20 °C.

## Structural basis of enhanced PA binding and reduced positional sensitivity to Ala mutation

To understand the molecular basis of improved binding, we attempted to solve the structures of P3, P4, P10 and P24 using different PA_Cter_ strains, either by testing co-crystallization using purified peptides, or by fusing the N-terminus of the peptides to the C-terminus of PA by a flexible (GS)_5_ linker. We obtained the crystal structures of A/California/07/2009 K369R-PA_Cter_ with P4, P10 and P24 bearing the flexible linker at 2.95–3.56 Å resolution (Supp. Table 1; Fig. 4). Each asymmetric unit contained three PA–peptide complexes related by non-crystallographic 3-fold symmetry (Supp. Fig. 3A). In all crystals, clear and continuous electron density defined a single peptide engaged in the canonical PB1-binding groove of PA_Cter_, adopting a 3_10_ structure like the wild-type (PDB 2ZNL) (16).

**Figure 4.**
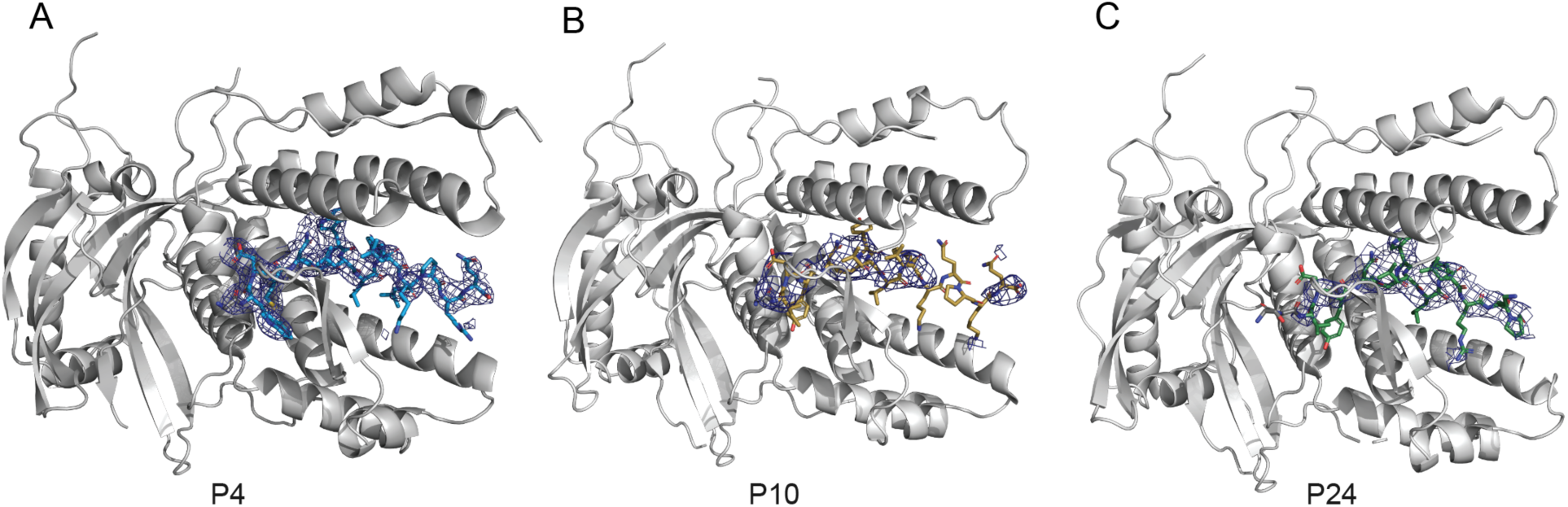
**Crystal structures of peptide–PA_Cter_ complexes**. (**A**) peptide 4 (P4), (**B**) peptide 24 (P24) and (**C**) peptide 10 (P10). Crystal structures depict the peptides bound within the PA groove, the N terminus on the left. Electron density is shown as a 2Fo–Fc map contoured at 1.0σ (blue mesh). Gray residues correspond to crystallographically resolved residues from the flexible linker. In peptide 24 (C) the last 3 C terminal residues are invisible. Detailed structural views are presented in Supp. Fig 3B.

Electron densities (2mFo–DFc maps contoured at 1.0σ) supported the placement of the bound helices in all three complexes. The P4 C terminus was defined in all three subunits (Fig. 4A) while the last three C-terminal residues were partially disordered in the high affinity P10 and P24 complexes (Fig. 4B, C). In P10, only one of the three subunits presented a visible density around position 15 at 1.0σ. No C-terminal density corresponding to the last three residues was visible for the remaining P10 subunits, as well as in all subunits of P24.

An alanine-scanning analysis, quantified by BLI on the WT peptide and P10 against WSN-PA_Cter_ demonstrated differences in positional sensitivity (Supp. Fig. 4). For WT, residues 4 to 10 were critical, as alanine substitution severely reduced the strength of interaction. In P10, only positions 4, 5 and 6 reduced the interaction severely. On the other hand, for P10, the C-terminal residues, mainly from positions 11 to 15, did not seem to affect binding, in accordance with the C-terminal flexibility observed in the crystal structure.

### Disruption of the PA-PB1 interaction and polymerase activity in cell-based assays

To assess whether the selected PB1-derived peptides could disrupt the PA–PB1 interaction from an influenza A virus (A/WSN/33) in living cells, we used a split-Gaussia luciferase complementation assay based on the transient expression of PA-Gluc1 and –PB1-Gluc2 constructs. Peptides of interest were expressed as peptide-EGFP constructs following plasmid transfection. To monitor the expression of the EGFP-fused peptides, EGFP fluorescence signals were measured using an Incucyte equipment, on HEK-293T transfected with the same transfection mixes as used for the split-luciferase assay. Peptide-EGFP levels for P3 and P4 were lower than WT, P10, and P24 in the PA-PB1 interaction assays (Supplementary Fig. 5A, B). There was no evidence of cytotoxicity associated with over-expression, as assessed with CellTiter-Glo viability assays (Supp. Fig. 5F). Normalized luminescence ratios (NLR), reflecting the PA-PB1 interaction, were determined as described previously (34). The WT peptide significantly reduced the interaction signal by 52.2% relative to the SCR peptide. The P3, P4, P10 and P24 peptides produced significantly stronger inhibition than the WT, with an 83–95% reduction of the interaction signal (P < 0.001; Fig. 5A). Pairwise comparisons showed that all four selected peptides significantly inhibited more strongly than WT, with no significant differences among the four (95% CI spanning zero; Supp. Table 4A). When the same assay was performed using the PA and PB1 proteins from an influenza B virus (B/Memphis/13/2003), the WT peptide had no inhibitory effect compared to SCR. P3, P4, P10, and P24 significantly reduced the interaction signal compared to SCR, although their effects were weaker compared to those observed with influenza A proteins (Fig. 5B; Supp. Table 4B). Overexpression of the GFP-fused peptides affected neither the unrelated Jun-Fos protein-protein interaction (Supp. Fig. 5E) nor the activity of the full-length Gaussia luciferase protein (Supp. Fig. 5F), supporting the specificity for the PB1-PA interaction.

**Figure 5.**
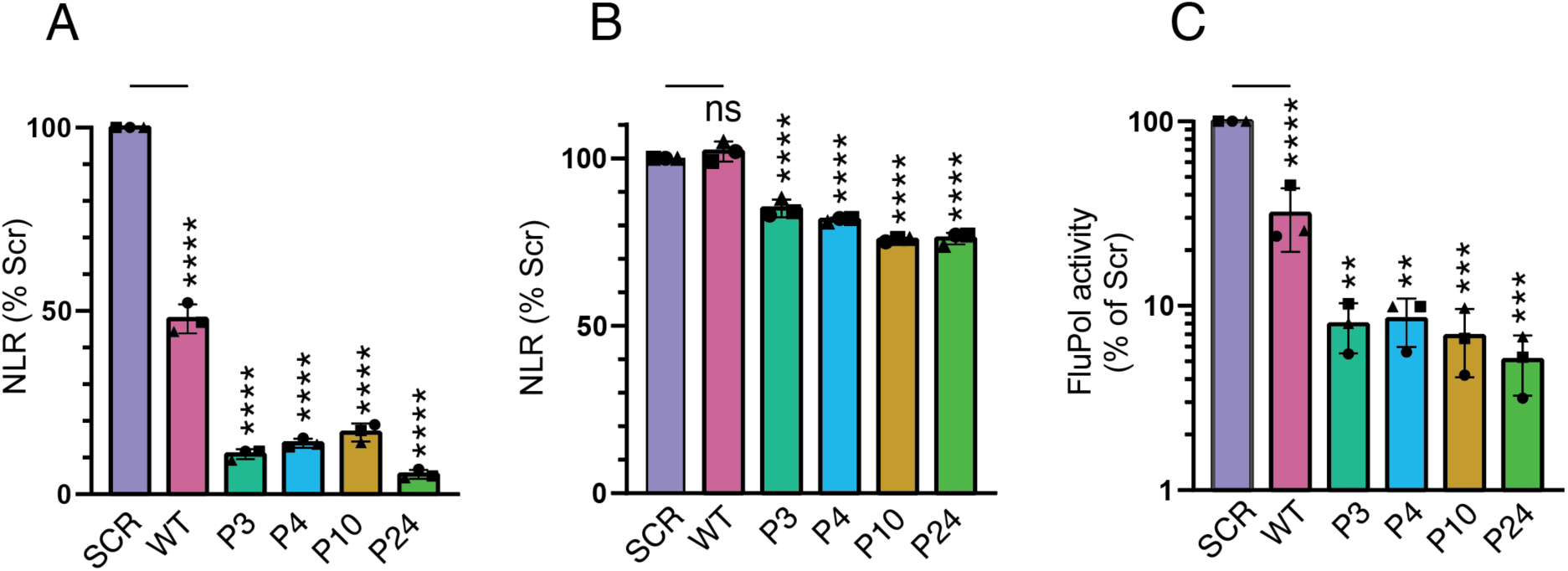
Phage display-selected PB1_1–15_-derived peptides disrupt influenza A polymerase complex formation and activity in cultured cells. (**A**, **B**) Split-luciferase assay measuring disruption of the PA-PB1 interaction from an influenza A (**A**) or influenza B virus (**B**). HEK-293T cells were co-transfected with plasmids encoding PB1 and PA fused to Gluc1 and Gluc2, respectively, together with expression plasmids encoding peptide-EGFP fusions. Active Gaussia luciferase is reconstituted when direct PA-PB1 interaction occurs. Luciferase activities were measured in cell lysates at 24 hours post-transfection (hpt) and normalized luminescence ratios (NLRs) were calculated and expressed as percentages relative to the SCR peptide. (**C**) Influenza A minireplicon assay. Viral ribonucleoproteins were transiently reconstituted in HEK-293T cells by transfecting expression plasmids for WSN-PB2, -PB1, -PA, -NP and a pseudo-viral RNA encoding the Firefly luciferase flanked by the 5’ and 3’ non-coding regions of the viral NS segment, in the absence or presence of the indicated peptide–EGFP constructs. As an internal transfection control, a pTK-Renilla plasmid was used. Luminescence was measured at 24 hpt. Firefly activity was normalized to Renilla activity and expressed as percentages relative to the SCR peptide condition. The data shown are the mean ± SD of three independent experiments performed in technical triplicates. Statistical significance relative to SCR is indicated: P < 0.01 (**), P < 0.001 (***) and P < 0.0001 (****); ns, not significant (two-way ANOVA; Tukey’s multiple comparisons test).

The peptides were further tested in a cell-based influenza A minireplicon assay. Transient co-expression of WSN-PB2, -PB1, -PA, -NP, together with a pseudo-viral reporter RNA encoding Firefly luciferase, was performed in the presence or absence of peptide–EGFP constructs, with Renilla luciferase as an internal control. The expression of the EGFP-fused peptides was monitored, and P3 and P4 were found to accumulate at lower levels (Supp. Fig. 5C), as in the previous PCA assays. The WT peptide reduced polymerase activity by 63.7% relative to the SCR peptide (p < 0.0001). Peptides P3, P4, P10, and P24 all produced a stronger inhibition, reducing polymerase activity by 92–95% compared with the SCR (p < 0.0001) and 75–80% compared to the WT (p < 0.01; Fig. 4C; Supp Table 3C). No significant differences were detected between the four selected peptides (Supp. Table 4C).

## Antiviral activity upon mRNA–LNP delivery

Preliminary plaque assays using synthetic peptides fused to the HIV-Tat, poly R (R8) and penetratin (YQIKIWFQNRRMKWKK) cell-penetrating peptides yielded unconclusive results, due to confounding toxicity effects (data not shown). Therefore, we used mRNA-lipid nanoparticle (mRNA-LNP) formulations encoding the peptide-GFP fusions as an alternative strategy for peptide delivery.

The physicochemical properties of the different mRNA-loaded LNP formulations (P10, P24, SCR, and WT fused with EGFP) were characterized in terms of particle size, polydispersity index (PDI), zeta potential, and mRNA encapsulation efficiency (Supp. Table 5). All formulations exhibited a similar hydrodynamic diameter ranging from 104.8 to 114.7 nm. The low PDI values (<0.12) indicated homogeneous particle size distribution. The surface charge of all LNPs was close to neutral, with zeta potential values ranging from −1.17 to 3.20 mV. All formulations showed high mRNA encapsulation efficiency, ranging from 83% to 92%.

Expression analyses, as monitored by GFP fluorescence signals (Fig 6A), showed that expression signals varied significantly with concentration only for WT and P10 between 0.1 and 1 µg of LNP per well (WT P = 0.0374; P10 P = 0.0337). Overall, a significant upward dose-expression trend was observed across the dataset (P = 0.0003), although P24 expression plateaued between 0.3 and 1 µg per well (P = 0.3939; non-significant). Significant overall difference between formulations were also observed (P = 0.0001); specifically, at 1 µg per well, P24 expression was significantly reduced compared to both SCR (P = 0.0176) and WT (P = 0.0128).

**Figure 6.**
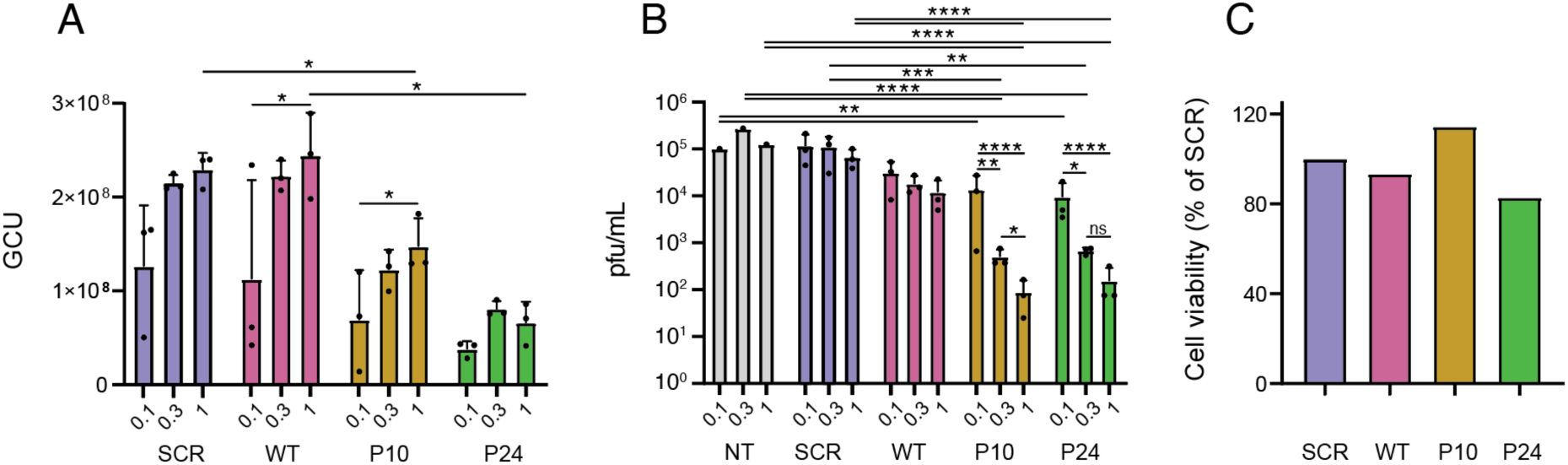
IAV antiviral activity of peptide-encoding mRNA-LNPs. **(A)** Expression of EGFP-fused peptides, as raw Green Calibrated Units (GCU), measured at 3 different doses (0.1, 0.3 and 1 µg per well) for SCR, WT, P10 and P24 mRNA-LNPs, n = 3 independent assays **(B)** Dose-dependent reduction in viral titers (PFU/mL) following treatment of A549 cells with mRNA-LNPs encoding GFP-fused SCR, WT, P10, or P24 peptides, at the indicated doses of 0.1, 0.3 or 1 µg per well of a 12-well plate and infection with the A/WSN/33 virus at a M.O.I of 0.001 PFU/cell. NT = no treatment control. n = 3 independent experiments; bars show mean ± SD. The data shown are the mean ± SD of three independent experiments performed in technical duplicates. **(C)** Cell viability assessed by CellTiter-Glo, expressed as % of untreated control cells, after treatment with mRNA–LNPs encoding SCR, WT, P10 or P24 at the highest dose of 1 µg per well. Significance levels as P < 0.05 (*), P < 0.01 (**), p < 0.001 (***) and P < 0.0001 (****) (Tukey’s multiple comparisons test).

The antiviral readout from plaque assays (Fig. 6B) following infection of 0.001 pfu/cell with the A/WSN/33 virus demonstrated significant main effects for both dose (P < 0.0001) and formulation (P < 0.0001). Within individual groups, SCR and WT showed no significant dose-dependent variation across the tested concentrations (P > 0.05), but both P10 and P24 exhibited strong, dose-dependent reductions in viral titers, decreasing significantly from 0.1 µg to 0.3 µg (P10: P = 0.0056; P24: P = 0.0106) and from 0.1 µg to 1 µg (P10: P < 0.0001; P24: P < 0.0001). Comparing formulations at each dose, SCR and WT viral levels were significantly higher than both P10 and P24 at 0.3 µg and 1 µg (P < 0.001), whereas at 0.1 µg, significant differences were primarily observed relative to SCR (vs. P10: P = 0.0064; vs. P24: P = 0.0005). The detailed information of the statistical analysis can be found in Supp. Table 6. There was no evidence of cytotoxicity at the highest dose, as assessed with CellTiterGlo assays (Fig. 6C), where all the tested conditions achieved more than 80% cell viability.

In summary, at the highest tested dose, P10 and P24 significantly reduced viral titers by ∼1,000-fold relative to the negative controls, and by ∼100-fold compared to the WT at the same dose.

## Discussion

In this study, we identified IAV PB1_1–15_-derived peptides that bind PA C-ter domain with higher affinity than the wild-type PB1_1–15_ peptide, disrupt polymerase assembly and activity in cell-based assays, and strongly inhibit IAV replication. We combined structure-guided library design with the diversity screening power of phage display and the analytical resolution provided by NGS to assess about a billion variants of the natural peptide, randomized at positions directly contacting PA. A previous mutagenesis and peptide array study mapped positional tolerances within the PB1_1–15_ peptide, providing a rational basis to design affinity enhanced peptides and assessed stepwise additive effects on affinity (35). By contrast, the library approach employed here allowed us to explore a greatly enlarged sequence space, allowing characterization of more complex mutation patterns and consequent non-linear effects on affinity. The findings indicate that the gains in affinity do not follow simple, position-specific rules. Instead, they may result from context-dependent combinations, which involve cooperative and compensatory mechanisms.

The PA_Cter_-peptide crystal structures show that the peptides adopt a 3_10_ conformation, in which their central hydrophobic cores appear resolved in the PA pocket, as demonstrated previously (16). However, the flanking regions also appear to play an important role in binding. In the wild-type structures, the first 15 amino acids of the PB1 N terminus were consistently resolved, even though longer PB1 constructs of 25 or 81 residues were used (15, 16). While the 10-residue PB1_2–11_ motif can bind stably (18), earlier observations indicated that flanking residues further stabilize the active peptide conformation (35). Accordingly, in crystal-resolved highest affinity peptides, P4, P10 and P24, the presence of an Arg or Lys at position 14 provides extra positive charge that may be involved in this interaction. P10 and P24 show no clear C-terminal density at 1σ. P24 has an extra charged residue (Lys) in position 12 whereas P10 has a Gln with a high hydrogen-bond potential and electron density was visible in one of the subunits. In contrast, P4 has an Leu in position 12, which effectively extends the hydrophobic core. This may explain why the C terminus of P4 is resolved, while the presence of polar or charged residues (such as Gln or Lys) may induce a more flexible, solvent-exposed state in P10 and P24. The lack of discrete hotspots on the C terminus in P10, together with the nanomolar affinities observed for P10 and P24, supports a model in which the less ordered C-terminal segment contributes to binding through distributed transient interactions rather than a single rigid interface (36, 37). Non-specific contacts and cooperative interactions may play a role, as observed elsewhere (38, 39).

Cellular assays support a direct competition model: in a split-luciferase-based protein complementation assay, co-expression of peptides blocked PA–PB1 heterodimer formation, while in minireplicon assays this translated into impaired FluPol activity (∼10-fold). This correlates with the biophysical data, where peptides that bind PA_Cter_ with higher affinity in vitro are better intracellular inhibitors of the PA-PB1 interaction.

A possible advantage of our peptide approach over small molecule inhibitors is that resistance-inducing escape mutations are likely constrained by the large binding interface area. The PA– PB1_1–15_ interface buries ∼1,000 Å², an order of magnitude larger than typical influenza small-molecule inhibitors (∼50–100 Å²). Binding is anchored by the conserved PB1 helical core within the PA pocket, where key contacts are distributed along the helix, characteristic of SLiM-mediated recognition (22, 23), while the terminal extensions appear to provide additional affinity through more distributed contributions. In this context, escape would likely require multiple coordinated substitutions rather than a single point mutation.

From a translational perspective, efficient vectorisation and intracellular delivery is a key objective. Our attempts to vectorize the lead peptides with appended TAT, penetratin and R8 sequences exhibited high toxicity in cellular assays and were discontinued (not shown). By contrast, encoding inhibitory peptides in mRNA-lipid nanoparticles (mRNA-LNPs) achieved robust intracellular expression as monitored through the GFP signal, with no detectable toxicity and a ∼1000-fold reduction in viral replication. This represents the first use of mRNA-LNPs for vectorization of antiviral peptides to our knowledge. This approach circumvents known limitations of intracellular-acting peptides such as cell-penetration, poor stability, rapid degradation (26, 29) and establishes a scalable, adaptable platform for other therapeutic targets.

Further in vivo studies will be necessary to determine efficacy, safe dosing, immune, and inflammatory effects of these LNPs upon intranasal administration. Nevertheless, recent formulation strategies have highlighted the potential of mucosal administration. Advances in thermoresponsive gelatin-PEG hydrogels for nasal mRNA delivery enable safe, sustained release and efficient transfection of airway epithelial cells (40). In parallel, maltodextrin-modified lipoplexes were shown to enhance mucus penetration and prolong nasal mRNA expression (41). Together, these advances provide a framework for adapting peptide-encoding mRNA formulations to intranasal or aerosol delivery against influenza and other respiratory infections.

By combining structure-based library design, high-throughput selection, functional validation, and LNP-based vectorization, we have demonstrated that PB1-derived linear peptides can strongly inhibit FluPol function in cellular assays of protein interaction, enzyme activity and infection. As such, they may open the way to a new class of anti influenza drug that has hitherto been held back by the challenges of peptide design and intracellular delivery.

## Methods

### PA_Cter_ expression and purification

PA_Cter_ (254–716, A/WSN/1933, UniProt P15659) was expressed as an N-terminal His–Smt fusion using SSGCID construct VCID7534. For phage display, a 6His–PA_Cter_–BAP construct was generated by AatII/NsiI insertion into pESPRIT002 (pET9a-derived). Expression in *E. coli* BL21-AI RIL (LB + 50 µg/mL kanamycin, 37 °C, 150 rpm) to OD_600_ 0.6–0.8, induced with 1 mM IPTG + 0.2% arabinose overnight at 18 °C. Harvested cells (4,500 rpm, 20 min, 4 °C) were lysed in Buffer 1 (25 mM Tris pH 8.0, 200 mM NaCl, 10 mM BME, 0.5% glycerol, 50 mM RE pH 8.0, 20% sucrose, lysozyme) for 30 min at RT and 30 min at 4 °C, then in Buffer 2 (as buffer 1, but with 25 mM imidazole, protease inhibitors), sonicated, and clarified (18,000 rpm, 45 min, 4 °C). Ni–NTA purification used 250 µL resin/L culture (30 min, 4 °C), washed to Bradford baseline, and eluted with 250 mM imidazole. Eluates were cleaved with ULP-1 (1:100 w/w, dialyzed overnight, 4 °C), reapplied to Ni–NTA, and flow-through (tag-free PA_Cter_) was purified by SEC on Superdex 200 (200 mM NaCl, 25 mM Tris pH 8.0, 1% glycerol, 10 mM BME, 50 mM RE pH 8.0). Fractions were assessed by SDS–PAGE, pooled, and stored at 4 °C.

### Phage Display Library Construction and Screening

FR (NNS at positions 1, 3, 4, 6, 11, 12, 14) and RR (DKS at 1, 3, 7, 8, 9, 10; NNS at 6) libraries with an N-terminal GSG linker were cloned into pADL-100 (Antibody Design Laboratories) via SfiI using an efficient library construction protocol (42). Libraries were electroporated into TG1 cells (50 ng or 1 µg DNA ×4), plated on 2×YT + 100 µg/mL ampicillin, and colonies were scraped and frozen. Phage were produced by growing library in LB to OD_600_ 0.4, superinfecting with 50 µL M13KO7d3 (Antibody Design Laboratories), incubating 30 min at 37 °C (static) + 30 min at 37 °C (180 rpm), inducing with 1 mM IPTG + kanamycin, and shaking overnight at 30 °C, 180 rpm. For panning, 50 µg biotinylated PA_Cter_ was immobilized on Dynabeads (Thermo Fisher Scientific) (1 mg R1, 0.1 mg R2, 0.01 mg R3) for 4 h at 4 °C, washed, blocked with SuperBlock (Thermo Fisher Scientific) (30 min, RT), and incubated with phage overnight. Beads were washed (R1: 6× PBS + 0.1% Tween-20 + 2× PBS; R2–3: 4× PBS + 0.1% Tween-20 + off-rate selection with 1 mg soluble PA for 2 h at 4 °C). Bound phages were eluted with 1 mL 1 M glycine-HCl (pH 2.0, 10 min), neutralized with 150 µL 1 M Tris-HCl (pH 9.0), and used to infect TG1 cells (OD600 0.4, 1 h, 37 °C).

### NGS and sequence analysis

DNA from 500 mL round 3 phage display scraped colonies was recovered using QIAprep Spin Miniprep Kit (Qiagen). Amplicons were generated via 25 PCR cycles (Phusion High-Fidelity DNA Polymerase; Thermo Fisher Scientific) with Illumina adapters (forward: 5’-ACACTCTTTCCCTACACGACGCTCTTCCGATCT-3’; reverse: 5’-GACTGGAGTTCAGACGTGTGCTCTTCCGATCT-3’) flanking a 500 bp phagemid region. Amplicons were purified (Macherey-Nagel NucleoSpin), normalized (500 ng at 20 ng/mL), and sequenced (Genewiz Amplicon-EZ). Sequences were trimmed to 15–110 bp (Phred >64), matched to the forward adaptor, translated, collapsed into unique peptides, and filtered at ≥5 observations. Pairwise Hamming distances were calculated using the R stringdist package, and peptides were partitioned into 20 sequence clusters.

### BioLayer Interferometry

Peptides were synthesized by SB Peptide (Grenoble) via SPPS and delivered crude in 100% DMSO or HPLC-purified, with QC by mass spectrometry. BLI experiments used OctetRED96e (Sartorius) with streptavidin sensors. Biotinylated peptides were immobilized to 1 nm and titrated with PA_Cter_ domain to establish optimal concentration range. Binding buffer: 200 mM NaCl, 25 mM Tris pH 8.0, 0.05% Tween-20. Protocol: 150 s baseline, 600 s association, 1500 s dissociation, 30 s regeneration in 1 M glycine-HCl (pH 2.0), 150 s wash. Sensors were reused. Kinetic constants (k_on_, k_off_, K_D_) were calculated using BLI software (Sartorius) assuming a 1:1 binding model.

### Thermal Shift Assay (TSA)

Thermal shift assays used a Stratagene MX100 RT-PCR instrument with 96-well optical plates. Reactions (20 µL) contained 2 µM PA_Cter_ protein, 10 µM peptide (WT, SCR, P3, P4, P10, P24), buffer (200 mM NaCl, 25 mM Tris pH 8.0, 50 mM TCEP), and SYPRO Orange (Thermo Fisher Scientific)(1:1000 dilution), with protein-only, peptide-only, and buffer-only controls. Samples were heated from 25 to 75 °C at 1 °C/min with fluorescence monitoring. Melting temperatures (T_M_) were derived from fluorescence first derivatives. Data were analyzed in R.

### Crystallization and structure resolution

Constructs fused (GS)_5_ linker + peptides to PA (A/California/07/2009, residues 257–716) via two-part overlap extension PCR using synthetic DNA cassettes (Thermo Fisher Scientific). P4 was cloned into VCID7534 (NcoI/XhoI); P10 and P24 were subcloned into P4 (SacII/NcoI). Constructs were amplified in *E. coli* Top10, expressed, and purified (PA_Cter_ protocol). Proteins (10 mg/mL) were screened at EMBL-HTX using CrystalDirect (43) with Wizard Classic I/II, Grid Screen Salt HT, and Morpheus matrices. Hits were optimized via hanging drop vapor diffusion. P4 crystals: 0.32 M NaH_2_PO_4_/0.68 M K_2_HPO_4_ (1 M, pH 7), cryo-protected with 10% then 20% glycerol. P10: 0.4 M NaH_2_PO_4_/0.76 M K_2_HPO_4_ (1.2 M, pH 6.9); P24: 0.32 M NaH_2_PO_4_/0.68 M K_2_HPO_4_ (1 M, pH 7), both cryo-protected with 20% glycerol. Crystals were flash-frozen in liquid nitrogen. X-ray data were collected at ESRF (ID23-2, ID30B/MASSIF-1) using MXCuBe (44), processed in EDNA (45), and solved via molecular replacement (MOLRP, CCP4, PDB: 2ZNL). Refinement used REFMAC5 (46) and COOT (47); model quality was assessed with SFcheck (48) and MolProbity (49).

### Cells and viruses

A549 (ATCC CCL-185) and HEK-293T (ATCC CRL-3216) cells were cultured in DMEM (Gibco) + 10% FCS + 1% penicillin-streptomycin. MDCK cells (National Reference Center for Respiratory Viruses, Institut Pasteur) were cultured in MEM + 5% FCS + 1% penicillin-streptomycin. Recombinant A/WSN/33 virus was produced from reverse genetics plasmids (gift from Pr. G. Brownlee, Oxford) (50). Reverse genetics efficiency was evaluated by supernatant titration on MDCK cells via plaque assay (51).

### Protein complementation assay

pCI-WSN-PB1-Gluc1, pCI-WSN-PA-Gluc2, pCI-Jun-Gluc1, pCI-Fos-Gluc2, and pCI-Gaussia-FL plasmids were used (34). GFP-fused peptide plasmids were generated by digesting pEGFP-N1 (GenBank U55762.1) with XhoI/BamHI, ligating with peptide-encoding oligonucleotides (45-nt), and transforming into *E. coli* Top10. HEK-293T cells (3 × 10^4^/well) were transfected with: 50 ng PA-Gluc1 + PB1-Gluc2 + 50 ng p-PB1-EGFP/empty pCI (PEI-max, 3 µL PEI at 1 mg/mL per 1 µg DNA); or B/Memphis PA/PB1-Gluc; or Jun/Fos-Gluc; or Gaussia-FL + empty pCI, all co-transfected with 50 ng p-PB1-EGFP or empty pCI. At 24 hpt, luciferase activity was measured (Centro XS LB960, 10 s reading after 50 µL Renilla reagent). Mean RLUs of triplicates and normalized luminescence ratios (NLR) were calculated. EGFP expression was monitored via Incucyte S3 (10×, 5 fields/well). Cell viability was assessed via CellTiter-Glo (Promega); toxicity was defined as <80% RLU vs. control.

### Polymerase activity assay

HEK-293T cells (3 × 10^4^/well) were co-transfected with 25 ng each pcDNA3.1-WSN-PB2, -PB1, -PA, -NP, plus 10 ng pPolI-Firefly and 5 ng pTK-Renilla (PEI). Additionally, 50 ng p-PB1-EGFP or empty pCI were co-transfected. Firefly and Renilla luciferase activities were measured via Dual-Glo system (Promega) at 24 hpt in technical triplicates.

### Formulation and characterization of mRNA-LNP

MM27 (O,O-dioleyl-N-histamine phosphoramidate, MW 692.05 Da) was produced in-house (52). β-sitosterol (MW 414.71 Da), DMG-PEG2000 (MW 2509.20 Da), and DOPE (MW 744.034 Da) were from Avanti Polar Lipids.

### mRNA

Cap1 poly(A) mRNA encoding WT, SCR, P24, or P10 fused to eGFP was produced by IVT. pT7 plasmids were linearized with SapI, and IVT performed using HiScribe T7 kit (Jena Biosciences) and dsRNA was removed via cellulose (53). Concentration and purity were determined by NanoDrop One and integrity and particle size using an Agilent 5200 Fragment analyzer.

### Microfluidic production of LNPs

LNPs were formulated using NanoAssemblr Ignite (Precision Nanosystems/Cytiva). Lipids (MM27:DOPE:β-sitosterol:DMG-PEG2000, 50:10:39:1) in ethanol were mixed with mRNA in 50 mM citrate (pH 4) at FRR 4:1 and TFR 11 mL/min, followed by 10-fold dilution in PBS and buffer exchange (Amicon Ultra-15, 100 kDa MWCO). LNPs were diluted in 10–30% sucrose (w/v) in PBS, frozen at -80°C, and thawed at RT for 30 min before use.

### Size, PDI and zeta potential

Hydrodynamic diameter and PDI were measured in DPBS via DLS (Zetasizer Ultra Red, Malvern). Zeta potential was measured in 10 mM NaCl by electrophoretic light scattering. Samples were diluted 500-fold.

### Encapsulation efficiency

mRNA encapsulation was assessed via RiboGreen assay. LNP samples and mRNA standards in TE buffer (with/without Triton X-100) were placed in black 96-well plates, RiboGreen dye added, and fluorescence measured (ClarioStar BMG, λex 485 nm, λem 528 nm). Encapsulation percentage = (fluorescence in TE)/(fluorescence in TE + Triton X-100) × 100, in triplicate.

### Viral inhibition assay

mRNA LNPs encoding peptides 10, 24, WT, or SCR were resuspended in PBS (5 mM). A549 cells (3 × 10^4/well) were infected with WSN (MOI 10^-4^ PFU/cell) for 1 h at 37°C in Opti-MEM media (Thermo Fisher Scientific) + 2% FCS. After virus removal, 100 µL medium with peptide concentrations was added in triplicate. Supernatants were pooled and virus production assessed via plaque assay (51). Cell viability was assessed via CellTiter-Glo assay.

## Acknowledgements

This project received funding from the Agence Nationale de Recherche (ANR2021 FluPept project, ANR-21-CE18-0024) and the Fondation pour la Recherche Médicale (FRM) (grant “Equipe 2017” DEQ20170336754). AFP received funding from the École Doctorale Chimie et Sciences du Vivant (EDCSV) – UGA. CP acknowledges funding from ANR France 2030 PEPR BBTI RNAvac.

The study used the platforms of the Grenoble Instruct-ERIC center (ISBG; UAR 3518 CNRS-CEA-UGA-EMBL) within the Grenoble Partnership for Structural Biology (PSB), supported by FRISBI (ANR-10-INBS-0005-02) and GRAL, financed within the University Grenoble Alpes graduate school (Écoles Universitaires de Recherche) CBH-EUR-GS (ANR-17-EURE-0003). The IBS acknowledges integration into the Interdisciplinary Research Institute of Grenoble (IRIG, CEA).

We acknowledge the European Synchrotron Radiation Facility for provision of beam time on ID23-2 and MASSIF-1 (ESRF BAG allocations MX2357 and MX2458). We thank Angeline Van Der Heyden from the Plateforme Caractérisation des Interactions at the Institut de Chimie Moléculaire de Grenoble. We thank Shibom Basu and Matthew Bowler for their support at ID23-2 and ID30-1/MASSIF-1 beamlines, respectively. We thank Khadija El Achi, Marah Saad and Olga Chirkova for technical contributions. We thank Stephen Cusack for the B/Mem plasmids, and Thomas Edwards and colleagues from the SSGCID for providing the expression vector VCID7534.

## Declaration of competing interests

AFP, WPM, NN, CI and DJH are inventors on patent applications filed by the CNRS on behalf of CNRS, UGA, CEA, IP and INSERM related to this work. The other authors declare no competing interests.

## Supplementary Figures

**Supplementary Figure 1.**
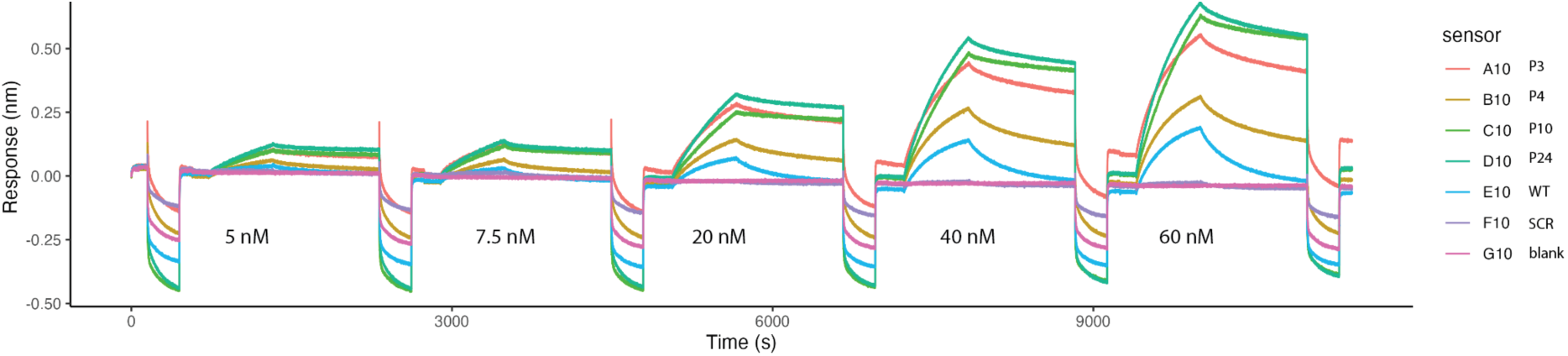
Raw BLI sensorgrams for peptide binding to PA_Cter_. Binding responses for peptides P3, P4, P10, P24, WT and SCR, and buffer blank measured at multiple PA_Cter_ concentrations (5, 7.5, 20, 40, 60 nM). Sensorgrams were collected on a Sartorius Octet RED96 with biotinylated peptides immobilized on streptavidin tips. Traces are shown as directly recorded, illustrating relative signal quality and specificity across peptides.

**Supplementary Figure 2.**
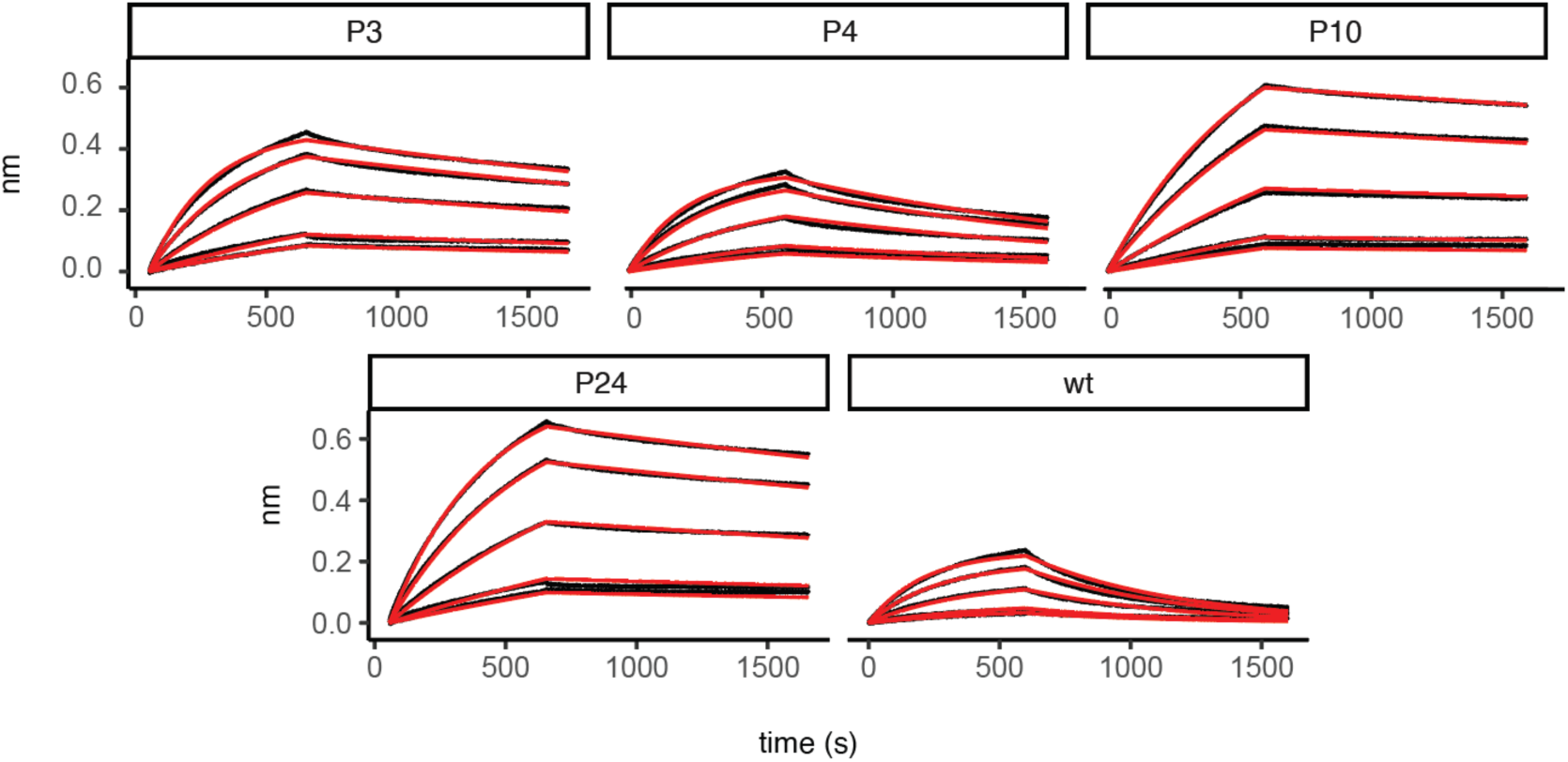
Global kinetic fits of BLI sensorgrams. Representative sensorgrams for peptides P3, P4, P10, P24 and WT (black) with corresponding global fits (red) overlaid. Kinetic parameters derived from these fits are reported in Fig. 2A and Table 1. Data shown are from a single side-by-side experiment performed under identical conditions to allow direct comparison of peptide variants.

**Supplementary Figure 3.**
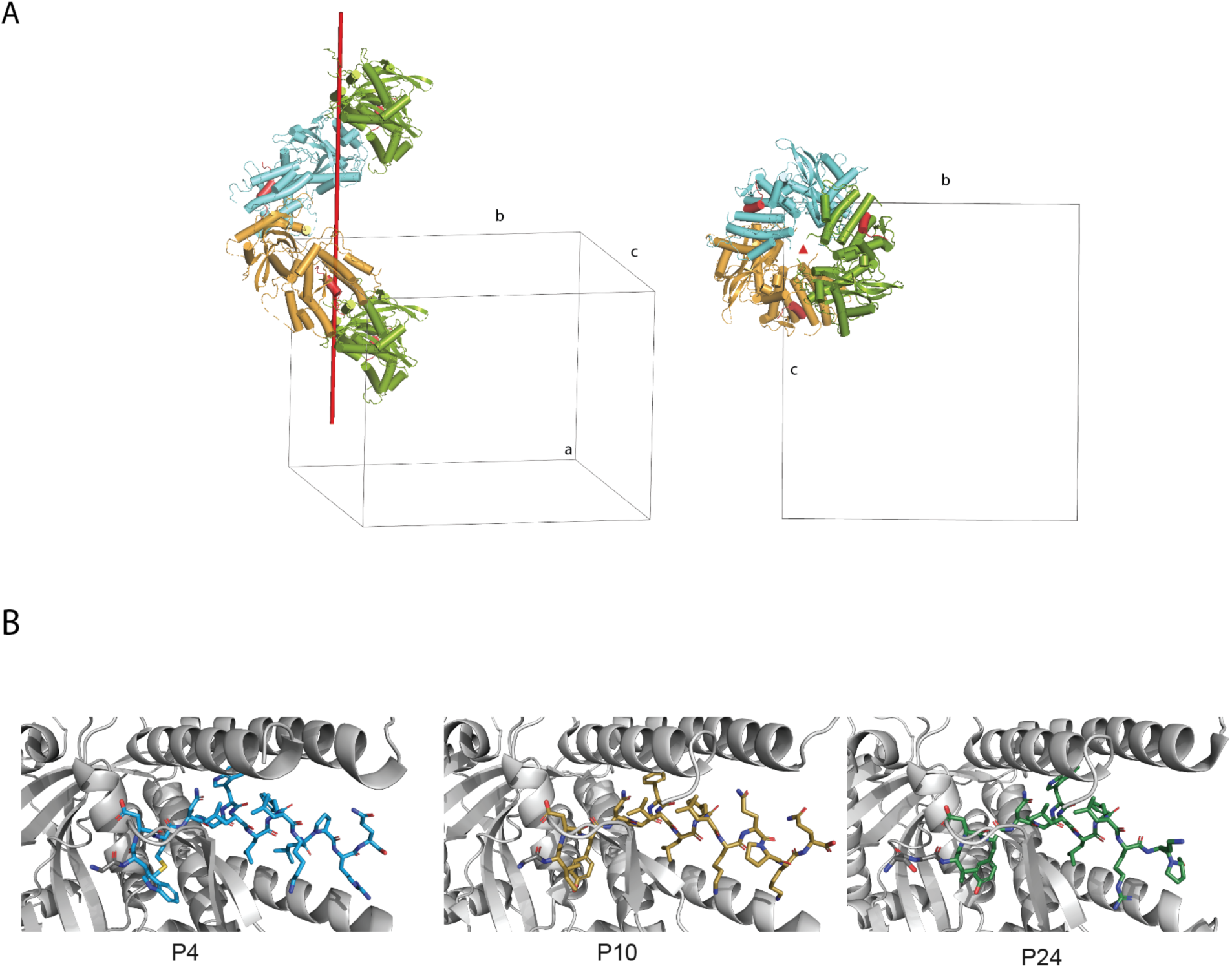
(**A**) The asymmetric unit (ASU) of the PA_Cter_–peptide crystal, showing three protomers per ASU (cylinder cartoon, colored orange, cyan and green), each with one bound peptide (highlighted in red). The red axis indicates a 3-fold non-crystallographic screw axis parallel to the ASU. **(B)** Close view of evolved peptides bound to PA_Cter_, from left to right P4, P10 and P24. The residues from the linker are colored gray.

**Supplementary Figure 4.**
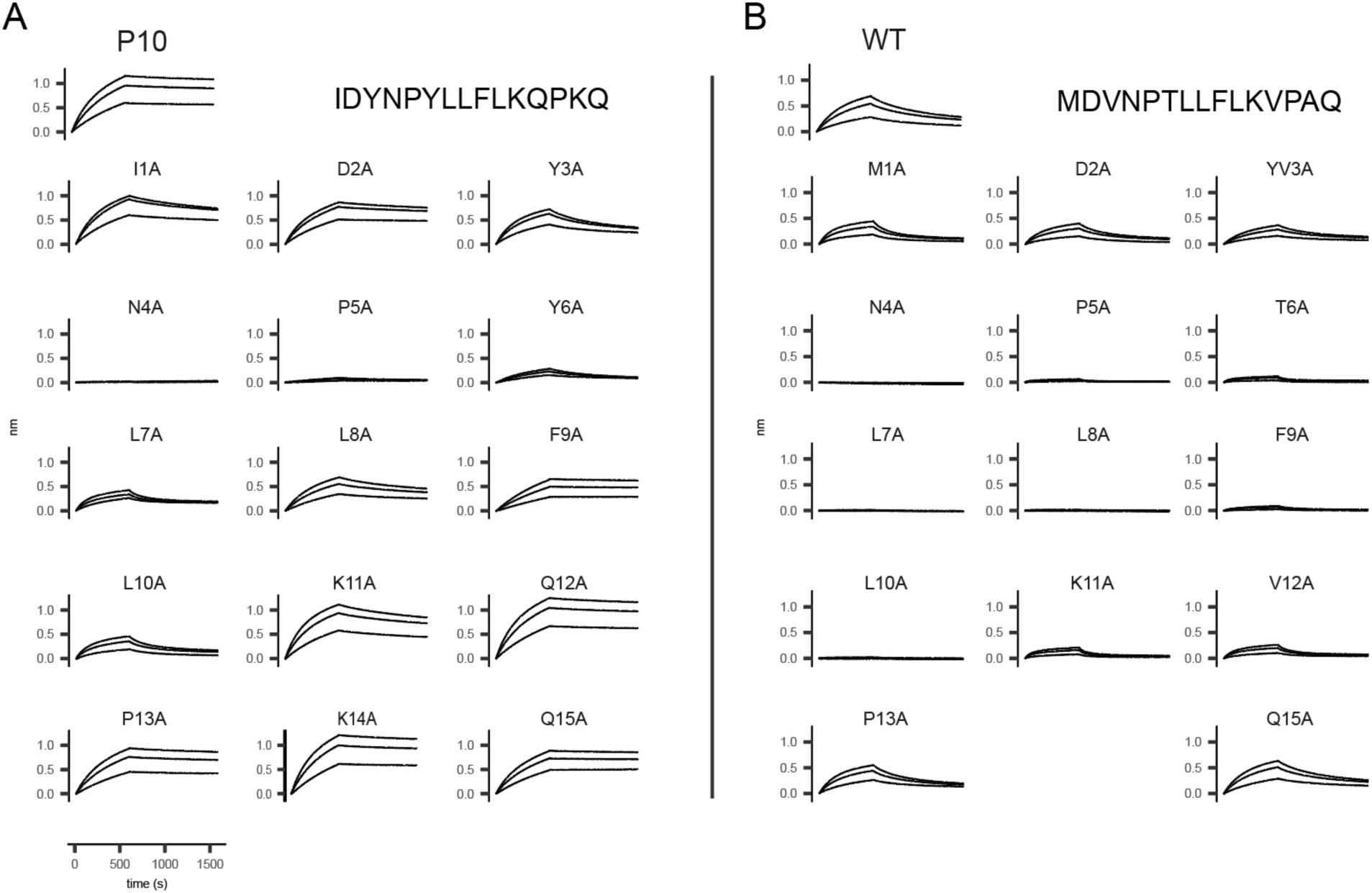
BLI sensorgrams from systematic alanine-scanning mutagenesis. (A) Peptide 10 and (B) wild type.

**Supplementary Figure 5.**
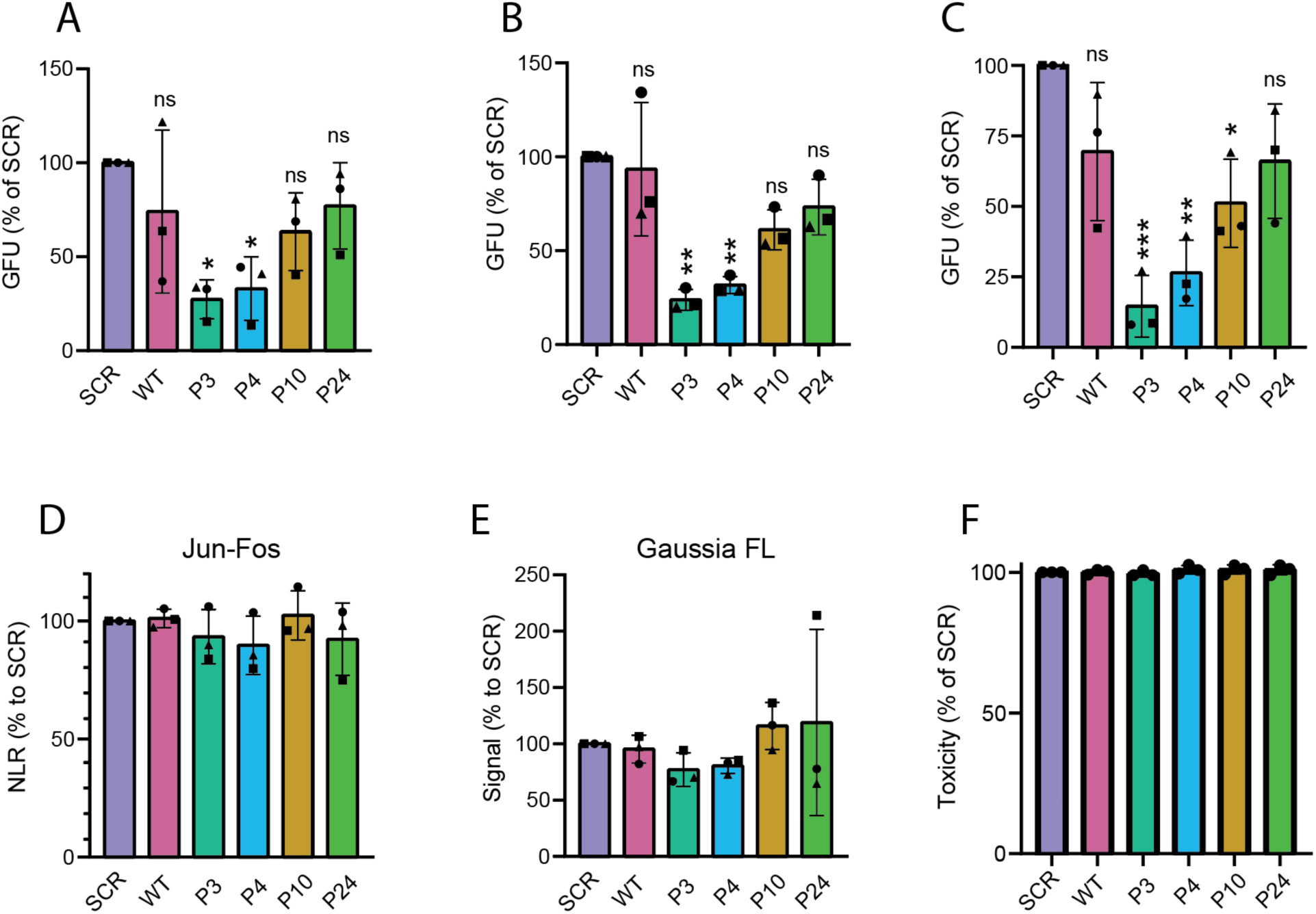
Expression, specificity and toxicity controls for EGFP-fused peptides. (A–C) Accumulation levels of the indicated EGFP-fused peptides as assessed by EGFP fluorescence signal in parallel with the PCA assay shown in Figure 4 (A, B) and the minireplicon assays shown in Figure 4 (C). (D) Split-luciferase complementation assay to assess the impact of EGFP-fused peptides on the Jun-Fos interaction. (E) Gaussia luciferase activity assay upon co-expression of the full-length Gaussia luciferase with the indicated GFP-fused peptides. (F) CellTiter-Glo viability assay upon expression of the indicated EGFP-fused peptides. Bars represent mean ± SD, individual data points, each corresponding to a biological replicate performed in technical triplicates (A-C) are shown; significance was determined by one-way ANOVA with Tukey’s multiple comparisons test. Statistical significance relative to SCR is indicated: P < 0.05 (*), P < 0.01 (**) and P < 0.001 (***); ns, not significant (Tukey’s multiple comparisons test).

## Supplementary Table

**Supplementary Table 1.** NGS library selection features.

|  | **Fully Randomized** | **Rationally Randomized** |
| --- | --- | --- |
| **DNA diversity** | $3.44 \times 10^{10}$ | $1.15 \times 10^9$ |
| **Protein Diversity** | $1.28 \times 10^9$ | $2 \times 10^8$ |
| **Library Size** | $7.7 \times 10^8$ | $7.1 \times 10^8$ |
| **Unique sequences** | 1,886 | 4,408 |
| **Total sequences** | 18,091 | 14,782 |

**Supplementary Table 2.**
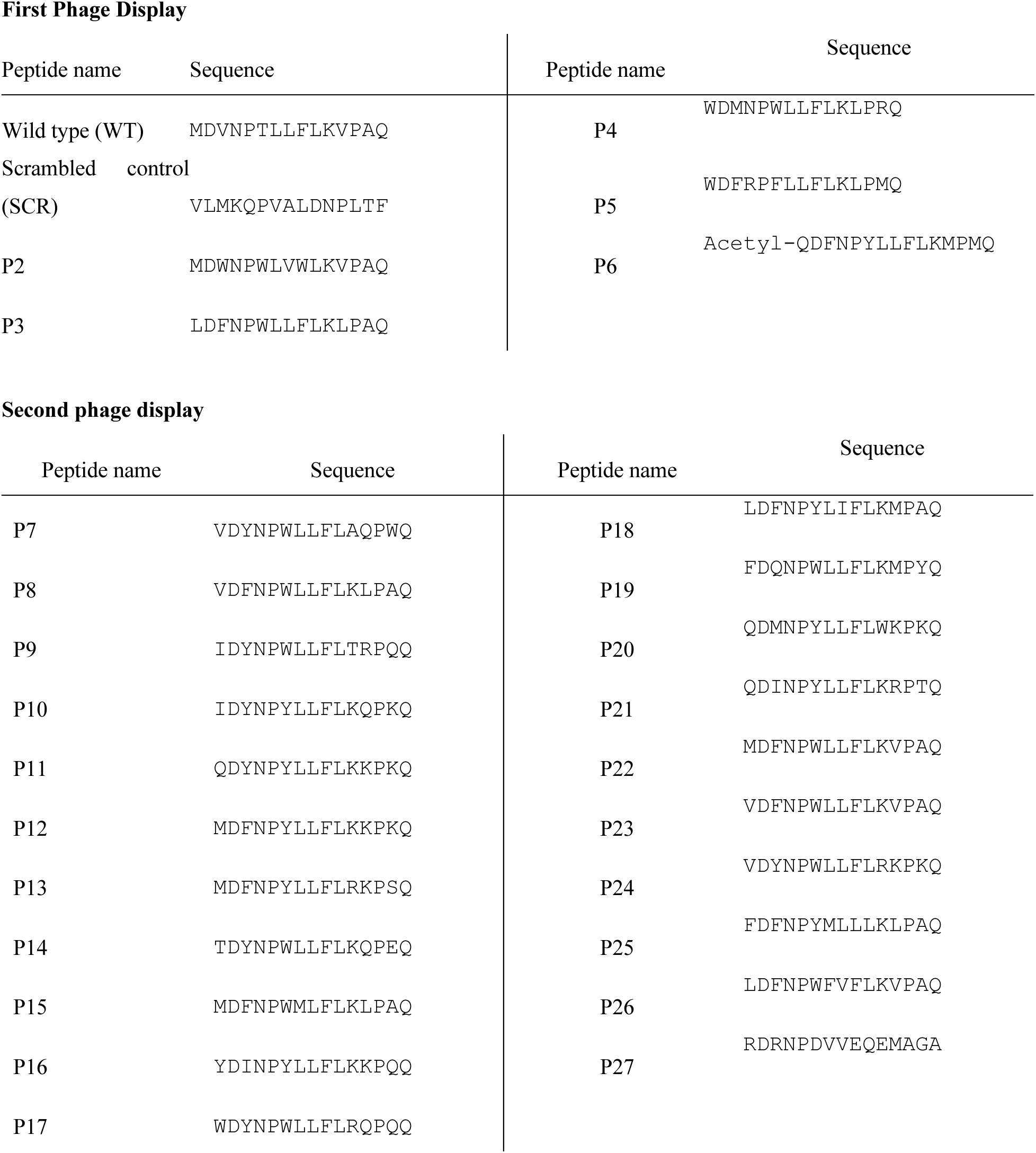
Initial peptide hit sequences as synthesized chemically for studies

**Supplementary Table 3.**
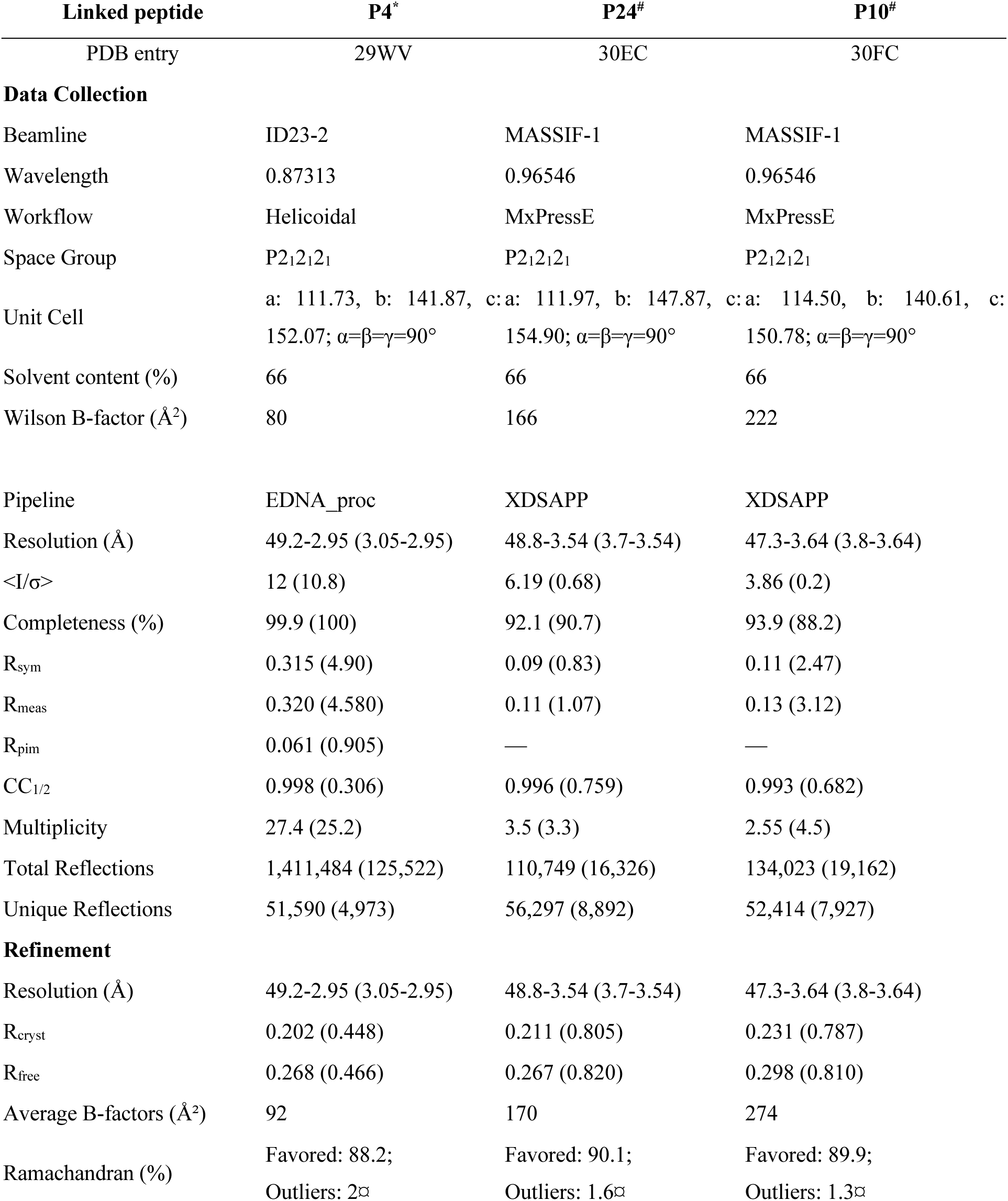
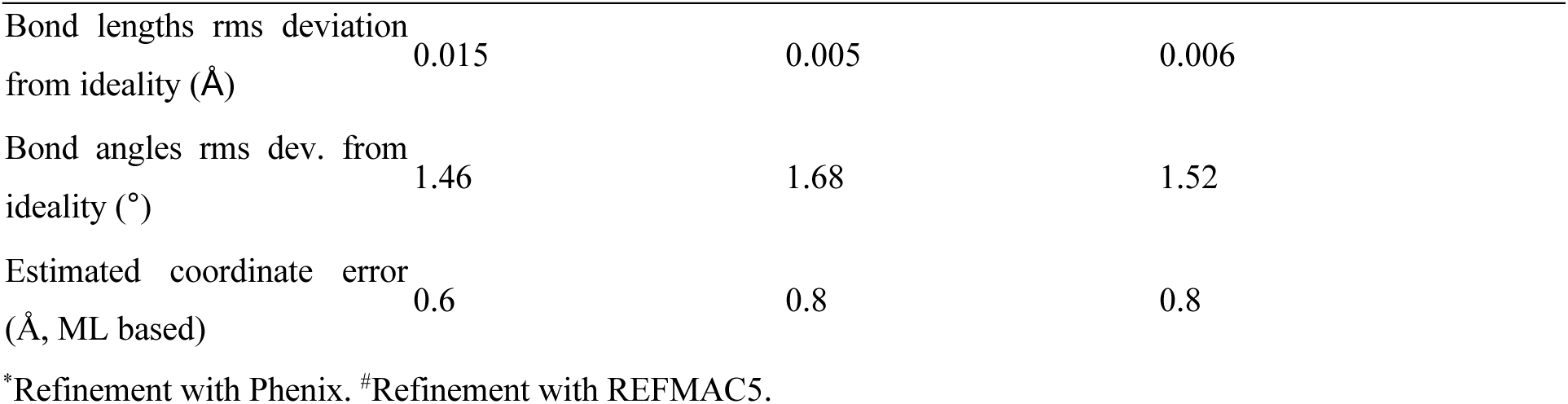
Data collection and refinement statistics for PA–peptide complexes. Strain: A/California/07/2009 K369R. Construct residues: 256–716

**Supplementary Table 4.** Tukey’s multiple comparisons test for cell-based assays of peptide inhibition. Pairwise of SCR, WT, and evolved peptides (P3, P4, P10, P24) in the PA–PB1 protein complementation assay (PCA, influenza A and influenza B) and the influenza A minireplicon assay. Mean differences, 95% confidence intervals (CI), Tukey adjusted P values, and significance summaries are reported.

**(A) PA–PB1 PCA, Influenza A (Fig. 5A)**
| Comparison | Mean Diff. | 95% CI of diff. | Significant? | Summary | Adjusted P value |
| --- | --- | --- | --- | --- | --- |
| SCR vs WT | 52.18 | 46.40 to 57.96 | Yes | **** | <0.0001 |
| SCR vs P3 | 89.10 | 83.32 to 94.88 | Yes | **** | <0.0001 |
| SCR vs P4 | 86.09 | 80.31 to 91.87 | Yes | **** | <0.0001 |
| SCR vs P10 | 83.19 | 77.42 to 88.97 | Yes | **** | <0.0001 |
| SCR vs P24 | 94.59 | 88.81 to 100.4 | Yes | **** | <0.0001 |
| WT vs P3 | 36.92 | 31.14 to 42.70 | Yes | **** | <0.0001 |
| WT vs P4 | 33.91 | 28.13 to 39.69 | Yes | **** | <0.0001 |
| WT vs P10 | 31.01 | 25.23 to 36.79 | Yes | **** | <0.0001 |
| WT vs P24 | 42.40 | 36.63 to 48.18 | Yes | **** | <0.0001 |
| P3 vs P4 | –3.01 | –8.79 to 2.77 | No | ns | 0.5289 |
| P3 vs P10 | –5.91 | –11.69 to –0.13 | Yes | * | 0.0441 |
| P3 vs P24 | 5.49 | –0.29 to 11.26 | No | ns | 0.0664 |
| P4 vs P10 | –2.90 | –8.68 to 2.88 | No | ns | 0.5644 |
| P4 vs P24 | 8.49 | 2.71 to 14.27 | Yes | ** | 0.0036 |
| P10 vs P24 | 11.39 | 5.61 to 17.17 | Yes | *** | 0.0003 |

**(B) PA–PB1 PCA, Influenza B (Fig. 4B)**
| Comparison | Mean Diff. | 95% CI of diff. | Significant? | Summary | Adjusted P Value |
| --- | --- | --- | --- | --- | --- |
| SCR vs WT | −2.00 | −6.97 to 2.97 | No | ns | 0.7522 |
| SCR vs P3 | 15.00 | 10.03 to 19.97 | Yes | **** | <0.0001 |
| SCR vs P4 | 18.33 | 13.37 to 23.30 | Yes | **** | <0.0001 |
| SCR vs P10 | 24.33 | 19.37 to 29.30 | Yes | **** | <0.0001 |
| SCR vs P24 | 24.00 | 19.03 to 28.97 | Yes | **** | <0.0001 |
| WT vs P3 | 17.00 | 12.03 to 21.97 | Yes | **** | <0.0001 |
| WT vs P4 | 20.33 | 15.37 to 25.30 | Yes | **** | <0.0001 |
| WT vs P10 | 26.33 | 21.37 to 31.30 | Yes | **** | <0.0001 |
| WT vs P24 | 26.00 | 21.03 to 30.97 | Yes | **** | <0.0001 |
| P3 vs P4 | 3.33 | −1.63 to 8.30 | No | ns | 0.2827 |
| P3 vs P10 | 9.33 | 4.37 to 14.30 | Yes | *** | 0.0004 |
| P3 vs P24 | 9.00 | 4.04 to 13.97 | Yes | *** | 0.0006 |
| P4 vs P10 | 6.00 | 1.04 to 10.97 | Yes | * | 0.0154 |
| P4 vs P24 | 5.67 | 0.70 to 10.63 | Yes | * | 0.0225 |
| P10 vs P24 | −0.33 | −5.30 to 4.63 | No | ns | 0.9999 |

**(C) PA–PB1 mini replicon, Influenza A (Fig. 5C)**
| Comparison | Mean Diff. | 95% CI of diff. | Significant? | Summary | Adjusted P Value |
| --- | --- | --- | --- | --- | --- |
| SCR vs WT | 68.49 | 54.17 to 82.80 | Yes | * | 0.0348 |
| SCR vs P3 | 92.07 | 77.76 to 106.4 | Yes | **** | <0.0001 |
| SCR vs P4 | 91.53 | 77.22 to 105.8 | Yes | **** | <0.0001 |
| SCR vs P10 | 93.15 | 78.83 to 107.5 | Yes | **** | <0.0001 |
| SCR vs P24 | 94.93 | 80.61 to 109.2 | Yes | **** | <0.0001 |
| WT vs P3 | 23.59 | 9.271 to 37.90 | Yes | **** | <0.0001 |
| WT vs P4 | 23.04 | 8.728 to 37.36 | Yes | ** | 0.0014 |
| WT vs P10 | 24.66 | 10.35 to 38.97 | Yes | ** | 0.0017 |
| WT vs P24 | 26.44 | 12.12 to 40.75 | Yes | *** | 0.0009 |
| P3 vs P4 | −0.54 | -14.86 to 13.77 | Yes | *** | 0.0005 |
| P3 vs P10 | 1.08 | -13.24 to 15.39 | No | ns | >0.9999 |
| P3 vs P24 | 2.85 | -11.46 to 17.17 | No | ns | 0.9998 |
| P4 vs P10 | 1.62 | -12.70 to 15.93 | No | ns | 0.9822 |
| P4 vs P24 | 3.40 | -10.92 to 17.71 | No | ns | 0.9987 |
| P10 vs P24 | 1.78 | -12.54 to 16.09 | No | ns | 0.9627 |

**Supplementary Table 5.** Physicochemical characterization of mRNA-loaded lipid nanoparticles (LNPs). Hydrodynamic diameter (size), polydispersity index (PDI), zeta potential, and mRNA encapsulation efficiency of the different LNP formulations (M-10, M-24, M-Scr, and M-WT). Data are presented as mean ± standard deviation (SD) of replicate measurements.

| **LNPs** | **Size (nm)** | **PDI** | **Zeta Potential (mV)** | **Encapsulation efficiency %** |
| --- | --- | --- | --- | --- |
| **P10** | 104.8 ± 1.78 | 0.044 ± 0.02 | 1.62 ± 0.71 | 90 |
| **P24** | 107.4 ± 0.93 | 0.042 ± 0.01 | -1.17 ± 1.79 | 91 |
| **SCR** | 106.3 ± 0.34 | 0.05 ± 0.01 | 3.2 ± 0.73 | 92 |
| **WT** | 114.7 ± 1.09 | 0.11 ± 0.03 | 1.45 ± 0.92 | 83 |

**Supplementary Table 6.**
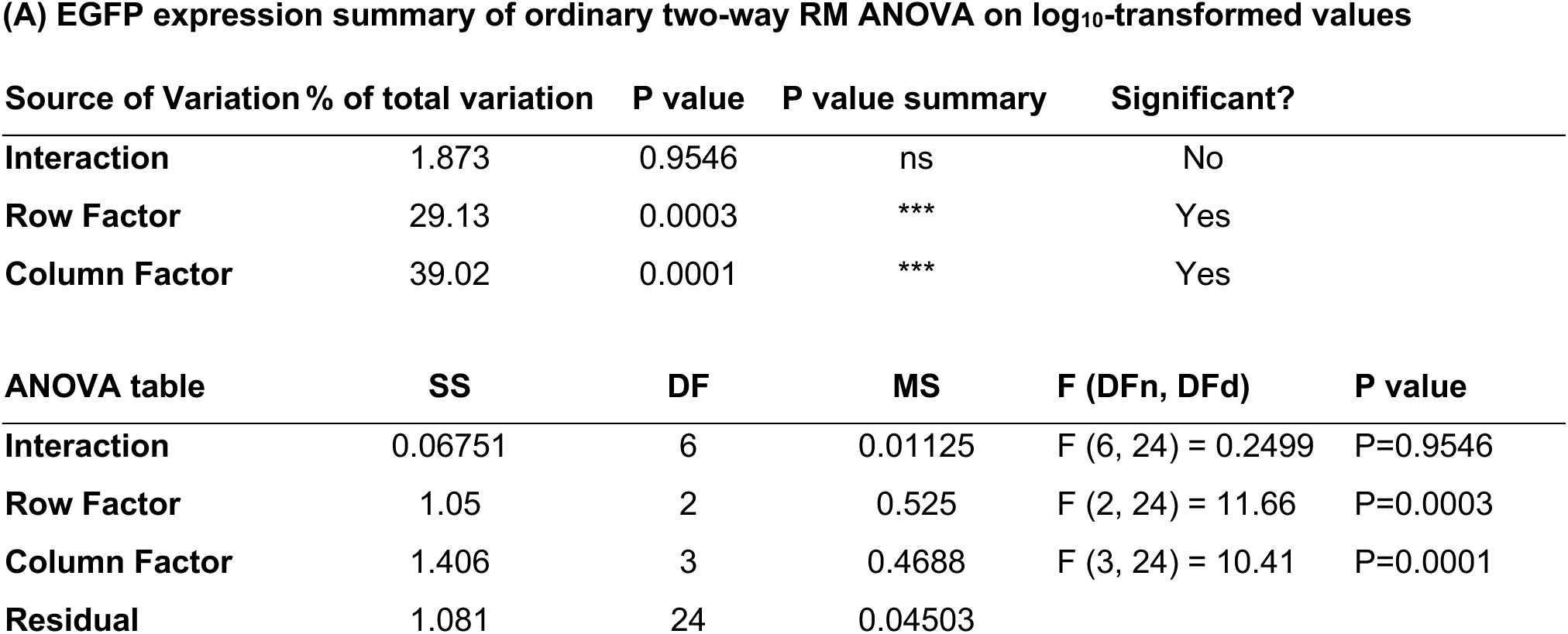
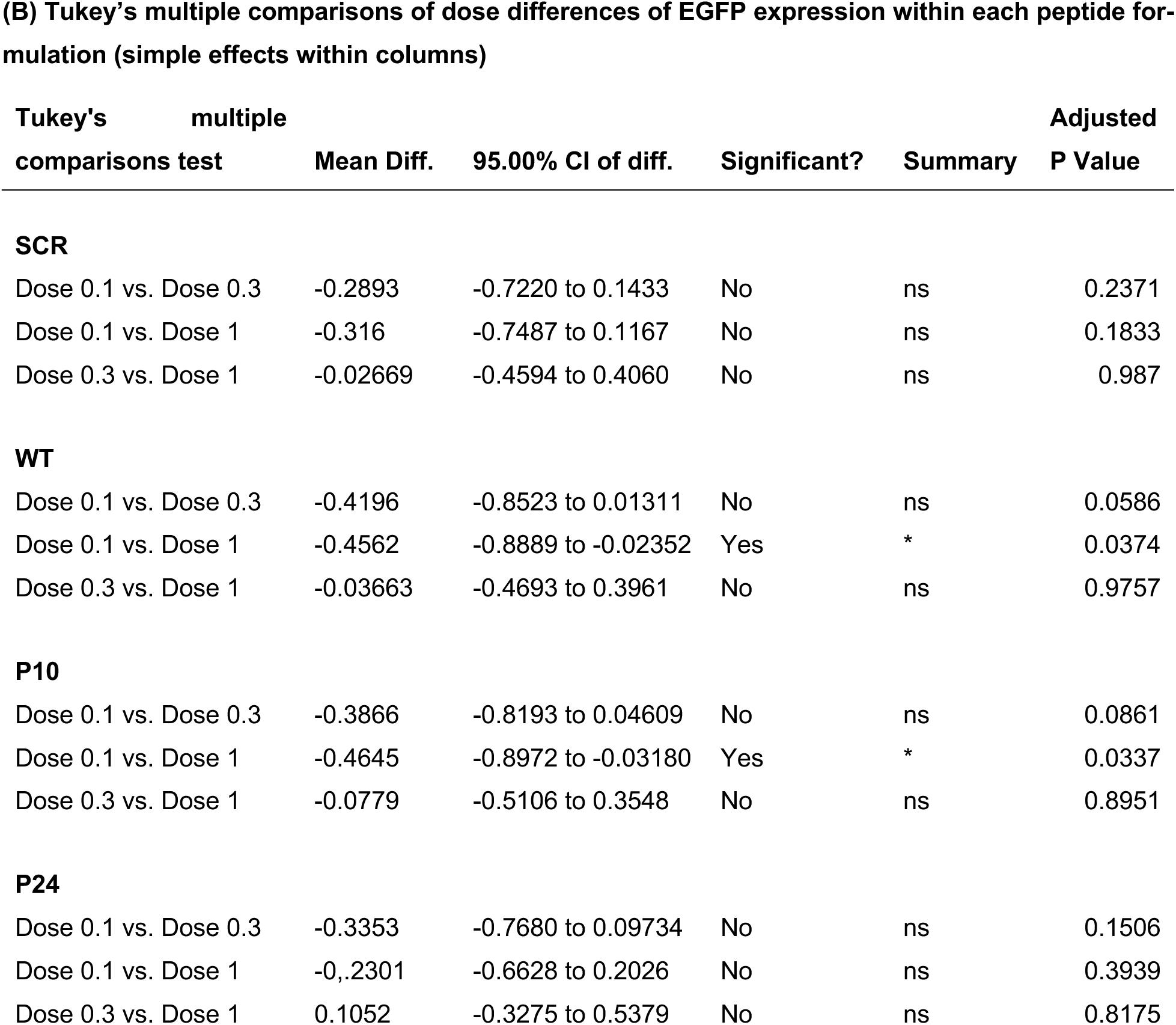
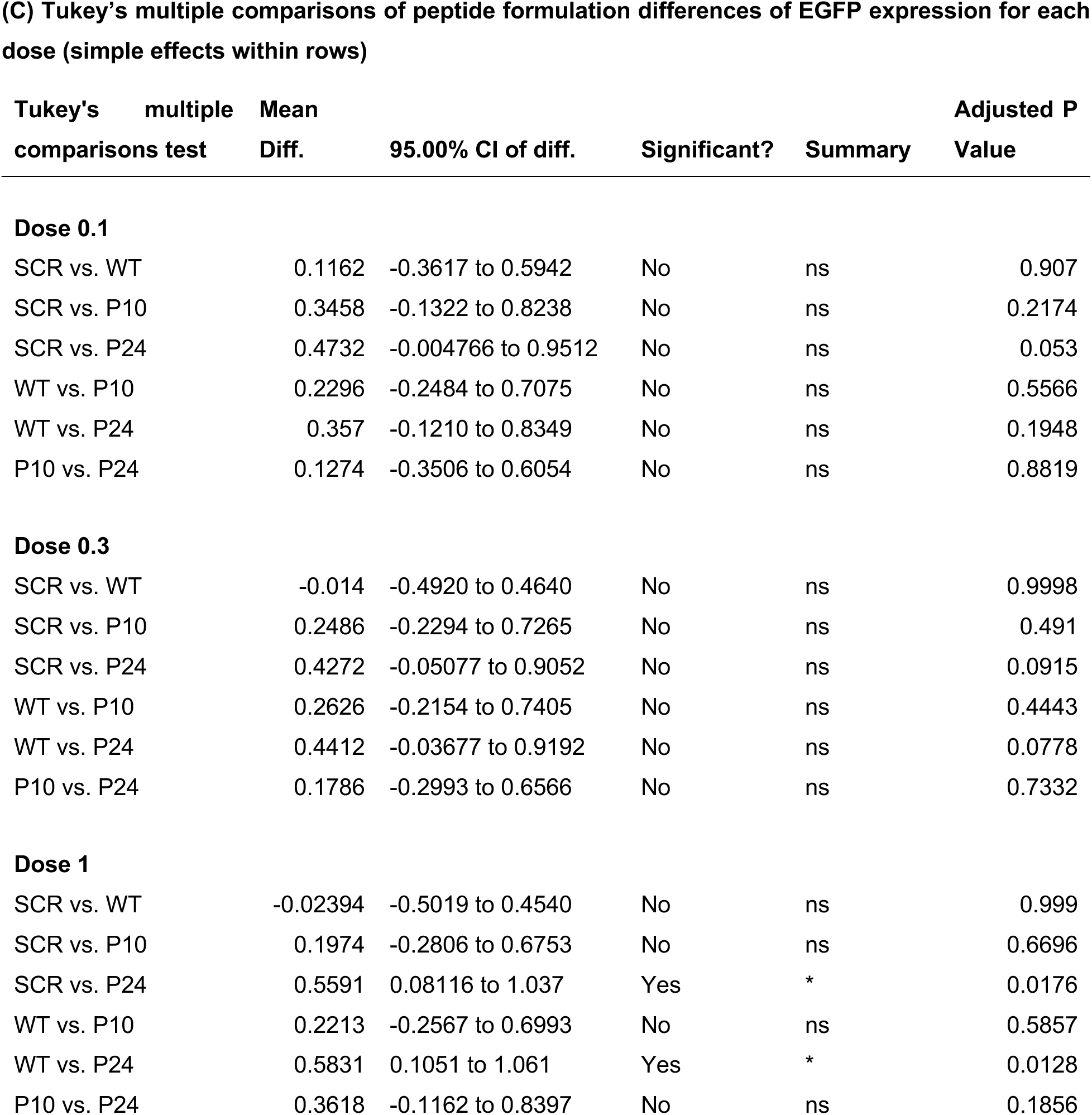
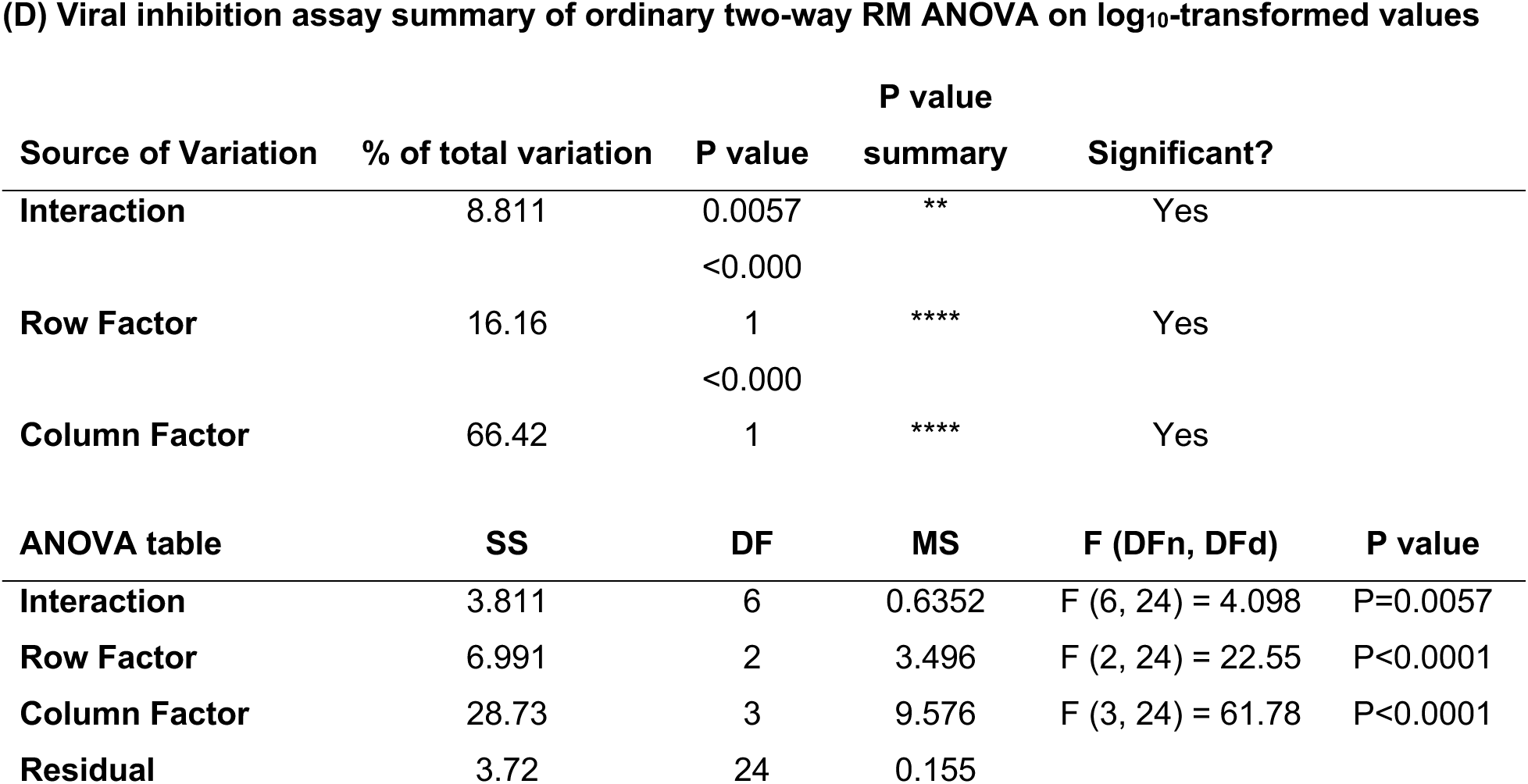
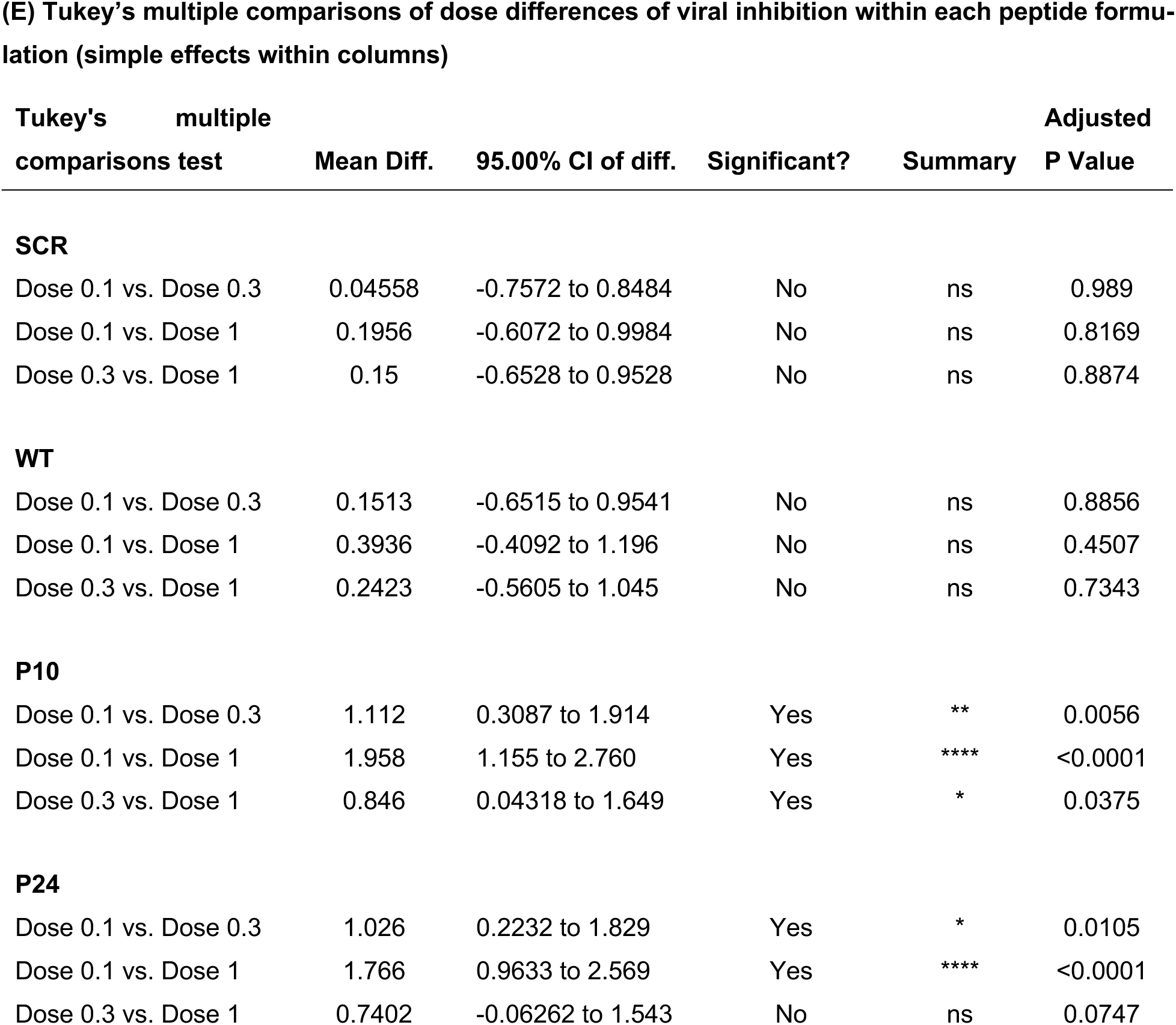
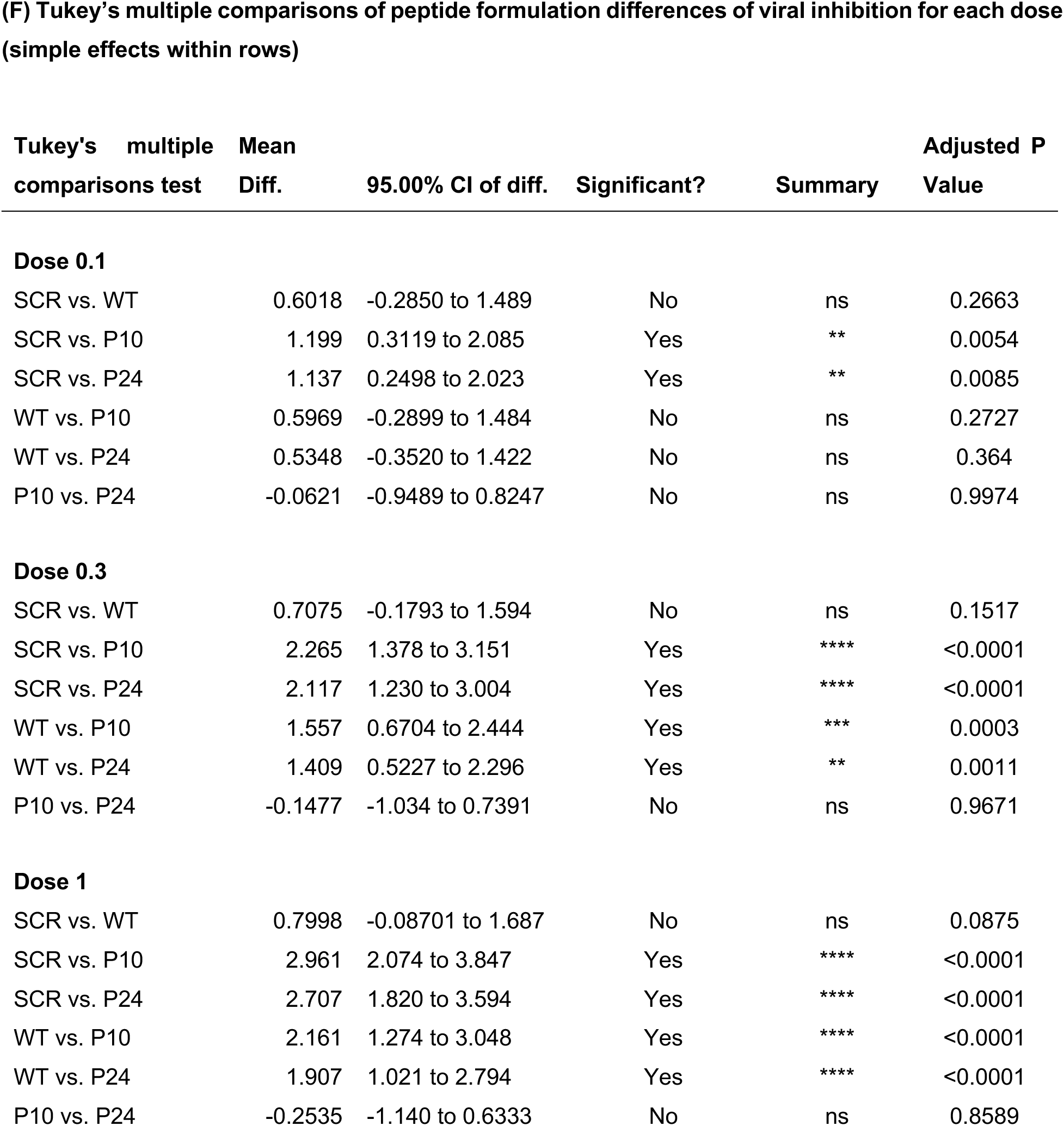
Two-Way RM ANOVA (analysis of variance) and post-hoc multiple comparisons analysis on log10-transformed for mRNA-LNP expression (A–C) and viral inhibition assays (D–F). **(A)** Two-Way RM ANOVA of EGFP expression vs mRNA-LNP doses (0.1, 0.3 and 1 μg per well). **(B)** Within-Column Multiple Comparisons (Comparing Doses Within Each Formulation). **(C)** Within-Row Multiple Comparisons of EGFP expression vs dose. (Comparing Formulations at Each Dose). **(D)** Two-Way RM ANOVA of plaque assay viral inhibition vs dose. **(E)** Within-Column Multiple Comparisons (Comparing Doses Within Each Formulation). **(F)** Within-Row Multiple Comparisons of EGFP expression vs dose. (Comparing Formulations at Each Dose). Mean differences, 95% confidence intervals (CI), Tukey-adjusted P values, and significance summaries are reported.

## Notes

### Summary of Updates

The title of the manuscript in the biorxiv metadata did not match exactly the title in the manuscript document. The uploaded manuscript file is identical to that previously submitted. We just updated the title in the metadata.

